# Spatiotemporal patterns of genetic diversity in the world’s coral reefs

**DOI:** 10.64898/2026.03.25.714244

**Authors:** Oliver Selmoni, Meredith C. Schuman

**Affiliations:** Remote Sensing Laboratories, Department of Geography, University of Zurich, Switzerland; Department of Chemistry, University of Zurich, Switzerland

**Keywords:** Coral reefs, macrogenetics, macrogenomics, climate change, genetic diversity, global biodiversity, framework

## Abstract

Coral reefs face widespread declines under global change, yet the status of their genetic diversity remains largely unknown. Here, we compiled genome-wide DNA sequencing data for 2,520 individuals from 18 reef taxa—including corals, fish, sharks, oysters, shrimp, sea anemones, and manta rays. These data were used to assess spatiotemporal patterns of genetic distances across 173 reefs worldwide between 1998-2018. While we did not observe an overall temporal decline in genetic distances, within-reef distances showed negative temporal trends, potentially reflecting population-level diversity loss. These effects varied across species but did not show clear distinctions between taxa. We then used satellite-derived seascape variables to predict local effects on genetic distances across reefs globally. Negative effects were predicted for the Red Sea, Northern Caribbean, and Coral Triangle, while positive effects were found across the South Pacific. Key predictors included declining oxygen levels, increasing nitrate concentrations, and rising water temperatures—variables that can be tracked in real time via Earth observation, enabling early warning for coral reef genetic diversity loss.

## 1. Intro

By signing the 2022 Kunming-Montreal Global Biodiversity Framework, most countries globally committed to protecting the genetic diversity of wild species and safeguarding their adaptive potential (*Kunming-Montreal Global biodiversity framework,* 2022). Monitoring is required to assess protection status, so there is an urgent need for tools that can establish a geographic baseline of genetic diversity and track its changes over time (Hoban et al., 2023). This monitoring task is particularly challenging in ocean ecosystems, where field-based DNA sampling is logistically complex (Planes and Allemand, 2023). Coral reefs exemplify this challenge: they are the most biodiverse ecosystems in our oceans and are undergoing a rapid degradation due to global change (Hughes et al., 2017), yet the status and trends of their genetic diversity remain largely unknown (Pinsky et al., 2023).

Macrogenetics is an emerging approach to quantify genetic diversity across species, ecosystems, and geographic regions (Leigh et al., 2021; Schmidt et al., 2023). This method typically involves compiling a database of genetic diversity metrics—such as nucleotide diversity or heterozygosity—from hundreds to thousands of previously published studies. These data are then jointly analyzed to identify large-scale patterns of variation across taxa, regions, and time. Applied to a wide range of organisms and ecosystems (Clark and Pinsky, 2024; Figuerola-Ferrando et al., 2023; Karachaliou et al., 2025; Leigh et al., 2019; Manel et al., 2020; Miraldo et al., 2016; Pelletier and Carstens, 2018; Schmidt et al., 2025, 2022, 2020; Shaw et al., 2025), the macrogenetic approach has led to important insights. For example, it enabled the estimation of a 6% global loss in genetic diversity across terrestrial and aquatic ecosystems since the Industrial Revolution (Leigh et al., 2019). In marine systems, macrogenetic studies have shown that fish genetic diversity correlates with environmental conditions (Clark and Pinsky, 2024; Manel et al., 2020) and anthropogenic pressures (Karachaliou et al., 2025), and that genetic diversity in habitat-forming species varies across taxonomic groups and along latitudinal gradients (Figuerola-Ferrando et al., 2023).

Despite its promise, the macrogenetic approach faces a sequence of challenges. First, genetic diversity metrics compiled from published studies can be noisy, as they are often based on heterogeneous sampling designs and analytical pipelines. This noise can mask true patterns of variation and reduce the ability to detect biologically meaningful trends (Exposito-Alonso, 2025). A promising solution is to bypass summary metrics and instead re-analyze raw genome-wide sequencing data from multiple previous studies using a standardized analytical pipeline (Exposito-Alonso et al., 2022). This approach offers greater power, as finer genomic resolution can reveal subtle differences in genetic structure between individuals (Leigh et al., 2021). A typical approach is the quantitative mapping of DNA sequencing reads against a reference genome (*i*.*e*, quantifying location and abundance of each DNA read from every individual). This mapping yields thousands of genetic variants that are then used to compare diversity between individuals (usually, biallelic single nucleotide polymorphisms, SNPs). However, this approach is more computationally demanding than the re-analysis of compiled genetic diversity metrics.

An innovative and, so far, rarely used option is k-mer-based analysis, which compares the frequency of short DNA substrings (k-mers) directly between DNA sequencing reads across individuals (Roberts et al., 2025). A key advantage is computational efficiency, as k-mer analysis does not require quantitative mapping of reads against a reference genome. Instead, k-mers require only a rapid mapping to remove contaminating sequences (Roberts et al., 2025), which can be achieved by checking whether the k-mers used to compare individuals are present in the reference genome sequence (or in a sequence database of non-target organisms) ((Kim et al., 2016)). This screening can be made even more efficient by randomly subsampling the set of k-mers used to compare individuals (Benoit et al., 2020; Fofanov et al., 2004). Another key advantage of k-mers is the broader representation of genetic variation. While mapping-based approaches often limit diversity analyses to biallelic SNPs, variation in k-mer frequencies can reflect a wide range of genetic differences (including SNPs, insertions, deletions, and translocations) (Roberts et al., 2025; Voichek and Weigel, 2020). A recent study on plants highlighted the relevance of this aspect: genetic diversity measured from k-mers has shown greater consistency with theoretical expectations from genome and population sizes, compared to traditional mapping-based methods (Roberts and Josephs, 2025).

A second major challenge in macrogenetic studies lies in the interpretation of observed spatiotemporal patterns of genetic diversity. Most studies attempt to explain these patterns by correlating genetic diversity with a limited set of predictors, such as geographic position (*e*.*g*, latitude) (Clark and Pinsky, 2024; Figuerola-Ferrando et al., 2023; Manel et al., 2020), environmental or bioclimatic conditions (*e*.*g*, mean annual sea surface temperature or chlorophyll concentration) ((Clark and Pinsky, 2024; Manel et al., 2020), or anthropogenic stressors (*e*.*g* human population density) ((Karachaliou et al., 2025). These predictors rarely explain more than 20% of the observed variation in genetic diversity. One likely reason is that commonly used variables may fail to capture the relevant ecological processes shaping genetic diversity (Leigh et al., 2021). For instance, sea surface temperature is often summarized using long-term means (Clark and Pinsky, 2024; Manel et al., 2020), but genetic diversity may also be shaped by temperature variability, seasonal anomalies, or recent trends in warming. Additionally, genetic diversity patterns are unlikely to be explained by single variables, and should instead be modeled considering the combined effect of a comprehensive set of predictors (Manel et al., 2020; Pelletier and Carstens, 2018). Identifying these key environmental predictors is essential for developing effective genetic monitoring systems, as many of these predictors can be tracked in near real-time through satellite Earth observation, offering a powerful early warning system to track conditions associated with genetic diversity loss (Schuman et al., 2023).

Here, we investigate how genetic diversity varies across space and time in the world’s coral reefs as revealed by the analysis of publicly available datasets (Figure 1). We used a k-mer approach to reprocess raw genome-wide DNA sequencing data from 18 reef species including fish, sharks, manta rays, corals, oysters, shrimps and sea anemones. This yielded individual-level genetic diversity estimates for 2,520 individuals sampled from 173 reefs globally (Figure 1A). We first assessed how genetic distances between individuals varied with taxonomy, geography, and year of sampling (Figure 1B). We then leveraged a comprehensive database of coral reef environmental trajectories (Selmoni et al., 2023) to identify the seascape conditions that best explained observed geographic patterns of genetic variation (Figure 1C-D), and finally used these relationships to predict genetic diversity across unsampled coral reefs worldwide (Figure 1E).

**Figure 1.**
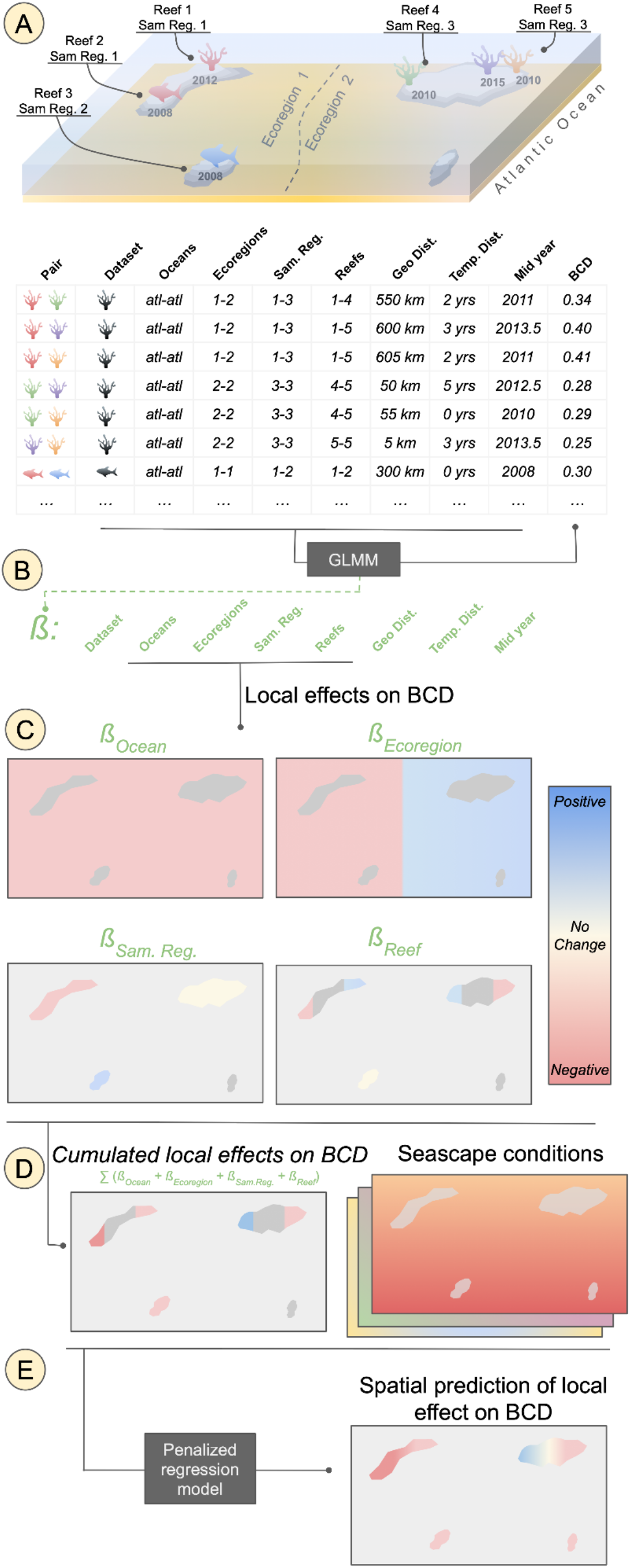
Modeling workflow. (A) Individual-level genomic data were compiled globally for various coral reef species, with known sampling site coordinates and years for each individual. Each sampling site was assigned to an ocean, ecoregion, sampling region (grouping sites within 100 km), and sampling reef (within 10 km). Bray-Curtis genetic distances (BCD) were then calculated between every pair of conspecific individuals. (B) A Generalized Linear Mixed Model (GLMM) assessed BCD variation across datasets, geographic regions, and sampling years. (C) The GLMM local effects on BCD were extracted for oceans, ecoregions, sampling regions, and sampling reefs. (D) For each sampling site, cumulative local effects on BCD (summing ocean, ecoregion, sampling region, and sampling reef effects) were calculated, alongside dozens of environmental variables characterizing seascape conditions. (E) A penalized regression model identified key seascape predictors of local BCD variation and generated spatial predictions for reefs worldwide.

## 2. Material and Methods

### Datasets selection and pre-processing of DNA sequencing reads

The datasets included in the analysis were selected through a manual search of peer-reviewed literature (August 2024). Using Web of Science and Google Scholar search functions, we first searched articles matching the keywords “coral reefs” and “population genomics”, which were then manually screened to find associated datasets that met the following criteria: (1) coral reef animal of any type, including habitat-building corals or reef-dwelling species such as fish, sharks or marine invertebrate; (2) individual genotyping of at least 50 individuals; (3) georeferenced sampling locations with reported sampling years; (4) at least two sampling locations (defined as group of samples collected up to 5 km apart); (5) a minimum spatial coverage of 50 km; and (6) genotyping performed using a restriction-site associated DNA sequencing (RAD-seq) method, with raw sequencing reads publicly available as short reads archived on NCBI.

For the 19 datasets matching the criteria (Table 1) (Bors et al., 2019; Buitrago-López et al., 2023; Gould and Dunlap, 2017; Lesturgie et al., 2023; Saenz-Agudelo et al., 2015; Salas et al., 2019; Selmoni et al., 2021; Shan et al., 2023; Sherman et al., 2020; Smith et al., 2022; Titus et al., 2024, 2019; Torquato et al., 2019; Walsh et al., 2022; Whitney et al., 2023), we downloaded raw sequencing reads from NCBI using a parallel implementation of the fastq-dump command from the sra-tools package (v. 3.1.1) (Leinonen et al., 2010). After an initial inspection of the read archives using FastQC (v. 0.12.1) (Andrews, 2010; Leinonen et al., 2010), we used TrimGalore (v. 0.6.10) (Martin, 2011) to filter low-quality reads, remove technical sequences (adapters and barcodes), and hard clip read ends—typically 2 nucleotides on the 3’ and 10–15 on the 5’ end (see Table S1 for dataset-specific cutoff values). Quality-filtered read archives were then re-evaluated with FastQC to ensure successful removal of technical sequence contamination.

**Table 1.** Genomic datasets. Overview of the 19 genomic datasets included in this study. For each dataset (row), the following information is provided: bibliographic reference (Ref.), species name, species type, ocean, ecoregion, number of samples before and after quality control (see Methods section on “Counting k-mer composition by sample”), number of sampling reefs (defined as sampling sites grouped within a 5 km radius), geographic range of the sampling reef distribution, and sampling period. For the total number of sampling sites, the number in parentheses indicates the number of unique sampling sites across all datasets.

| # | Ref. | Species | Type | Ocean | Ecoregions | # samples |  | # of sam. reefs | Geo. range [km] | period |
| --- | --- | --- | --- | --- | --- | --- | --- | --- | --- | --- |
|  |  |  |  |  |  | b. QC | a. QC |  |  |  |
| 1 | (Titus et al., 2024) | *Ancylomenes pedersoni* | Shrimp | Atlantic | Bahamian,Eastern Caribbean,Western Caribbean,Southern Caribbean,Floridian,Southwestern Caribbean,Bermuda | 148 | 132 | 14 | 3137 | 2013-2017 |
| 2 | (Buitrago-López et al., 2023) | *Stylophora pistillata* | Coral | Indian | Northern and Central Red Sea,Southern Red Sea | 448 | 393 | 11 | 1531 | 2011-2012 |
| 3 | (Buitrago-López et al., 2023) | *Pocillopora verrucosa* | Coral | Indian | Northern and Central Red Sea,Southern Red Sea | 316 | 259 | 10 | 1526 | 2011-2012 |
| 4 | (Smith et al., 2022) | *Platygyra daedalea* | Coral | Indian | Arabian (Persian) Gulf,Gulf of Oman | 170 | 148 | 15 | 1141 | 2013 |
| 5 | (Selmoni et al., 2021) | *Pocillopora damicornis* | Coral | Pacific | New Caledonia | 128 | 124 | 16 | 396 | 2018 |
| 6 | (Selmoni et al., 2021) | *Pocillopora acuta* | Coral | Pacific | New Caledonia | 150 | 133 | 16 | 396 | 2018 |
| 7 | (Selmoni et al., 2021) | *Acropora millepora* | Coral | Pacific | New Caledonia | 187 | 170 | 18 | 402 | 2018 |
| 8 | (Devlin-Durante and Baums, 2017) | *Acropora palmata* | Coral | Atlantic | Bahamian,Floridian,Greater Antilles,Eastern Caribbean | 72 | 63 | 11 | 1818 | 2002-2010 |
| 9 | (Titus et al., 2019) | *Bartholomea annulata* | Sea Anemone | Atlantic | Bermuda,Bahamian,Eastern Caribbean,Western Caribbean,Floridian,Southwestern Caribbean | 122 | 72 | 14 | 3060 | 2013-2018 |
| 10 | (Shan et al., 2023) | *Pinctada fucata* | Oyster | Pacific | Gulf of Tonkin,South China Sea Oceanic Islands | 74 | 69 | 4 | 1517 | 2017-2018 |
| 11 | (Whitney et al., 2023) | *Mobula alfredi* | Manta | Pacific | Hawaii | 37 | 35 | 3 | 152 | 2009-2015 |
| 12 | (Torquato et al., 2019) | *Pomacanthus maculosus* | Fish | Indian | Arabian (Persian) Gulf,Gulf of Oman,Western Arabian Sea,Gulf of Aden,Southern Red Sea,Northern and Central Red Sea,Northern Monsoon Current Coast | 151 | 149 | 23 | 3485 | 2012-2017 |
| 13 | (Bors et al., 2019) | *Pterois volitans* | Fish | Atlantic | Bahamian,Western Caribbean,Greater Antilles,Floridian,Eastern Caribbean | 120 | 104 | 21 | 2547 | 2012-2016 |
| 14 | (Salas et al., 2019) | *Dascyllus trimaculatus* | Fish | Indian | Northern and Central Red Sea,Gulf of Aden,Arabian (Persian) Gulf,Chagos,East African Coral Coast,Western and Northern Madagascar | 93 | 93 | 12 | 5729 | 1998-2013 |
| 15 | (Gould and Dunlap, 2017) | *Siphamia tubifer* | Fish | Pacific | South Kuroshio | 280 | 195 | 8 | 155 | 2012-2014 |
| 16 | (Saenz-Agudelo et al., 2015) | *Amphiprion bicinctus* | Fish | Indian | Gulf of Aden,Southern Red Sea,Northern and Central Red Sea | 103 | 62 | 10 | 2647 | 2009-2013 |
| 17 | (Sherman et al., 2020) | *Epinephelus striatus* | Fish | Atlantic | Bahamian | 96 | 34 | 7 | 579 | 2017 |
| 18 | (Lesturgie et al., 2023) | *Carcharhinus amblyrhynchos* | Shark | Pacific,Indian | New Caledonia,Western and Northern Madagascar,Line Islands,Phoenix/Tokelau/Northern Cook Islands,Tuamotus,Society Islands | 173 | 137 | 17 | 17046 | 2017 |
| 19 | (Walsh et al., 2022) | *Carcharhinus amblyrhynchos* | Shark | Indian, Pacific | Chagos,Cocos-Keeling/Christmas Island,Coral Sea,Papua,Ningaloo,Torres Strait Northern Great Barrier Reef,Central and Southern Great Barrier Reef,Exmouth to Broome | 169 | 148 | 9 | 8762 | 2013 |
| **TOTAL** |  |  |  |  |  | **3037** | **2520** | **239 (173)** |  | **1998-2018** |

Unbalanced read numbers and lengths between samples can introduce bias in downstream comparisons of k-mer counts (Hrytsenko et al., 2025). Additionally, processing large volumes of sequencing reads from thousands of samples across multiple datasets can be computationally intensive. To address these issues, we subsampled the same number of reads per sample within each dataset. Specifically, for each dataset, we sampled a number of reads corresponding to the 10th percentile of the read count distribution across all samples (Figure S1). This downsampling was not expected to affect downstream analyses, as we checked in one dataset with extensive sample size (grey reef shark from Indo-Pacific(Walsh et al., 2022); Figure S2). We also trimmed all reads to a fixed length equal to the median read length for that dataset (Table S1). These steps were performed using seqtk (v. 1.4) (Li, 2025).

### Counting k-mer composition by sample

We counted k-mers in each dataset using kmc (v. 3.2.4) (Kokot et al., 2017). Following previous studies using k-mers to assess population structure and genetic diversity, we counted canonical k-mers (*i*.*e*., each k-mer and its reverse complement were treated as the same) of length 31. Rare k-mers (counts < 5) were discarded, as they potentially result from sequencing artifacts. To optimize computational efficiency and standardize processing across datasets (see also previous section), we randomly sampled 500,000 k-mers per dataset and recorded their counts in a k-mer-by-sample matrix. This sampling was done using a custom R script (available in the online material).

As some k-mers may originate from non-target DNA (*e*.*g*., bacterial or other species contamination), we filtered the k-mer-by-sample matrix to exclude samples potentially enriched with non-target DNA. This was done by checking FastQC GC content in the sequencing reads of every sample and by excluding outlier samples (*i*.*e*., those with GC content outside the lower/upper quartile ± the interquartile range, IQR). Furthermore, we filtered out k-mers that might originate from non-target DNA. For this, two alternative filtering strategies were used to remove potential contaminants. For species with a reference genome available for a closely related species (within the same genus; 16 out of 19 datasets; Genbank identifiers in Table S1), we mapped k-mers against the reference genome using NextGenMap (v. 0.5.5) (Sedlazeck et al., 2013) and discarded any unmapped k-mers using Samtools (v. 1.16.1) (Li et al., 2009). For species lacking a suitable reference genome (3 out of 19 datasets), we mapped k-mers against the compressed database of Bacteria, Archaea, Viruses, and Human genomes of Centrifuge (v. 1.0.4.2) (Kim et al., 2016) and discarded any k-mers matching this database. We preferred reference-based filtering because it enables more specific exclusion of contaminant k-mers, including those originating from other animals (*e*.*g*., environmental DNA common in aquatic samples) or symbiotic organisms (which is particularly relevant for coral species) (Maire et al., 2021). As a quality control step, we compared the GC content in the k-mers sequences before and after this taxonomic filtering, under the assumption that contaminant reads would increase variability in GC content (Figure S3).

We finally refined each dataset’s k-mer count matrix, retaining only k-mers present in at least 5% of all samples. For every dataset, numbers of kmers and samples after each of these filtering steps are reported in Table S1.

### K-mers to characterize genetic variation within datasets

For each dataset, we normalized the k-mer count matrix using the Trimmed Mean of M-values normalization method (TMM; R package edgeR, v. 4.0.16) ((Robinson et al., 2009). TMM corrects compositional and library-size bias in counts from sequencing data and is more robust than standard normalization, because it reduces the influence of k-mers with extreme counts when estimating library sizes.

As in previous work using k-mers to measure genetic distances (Roberts et al., 2025; Roberts and Josephs, 2025), we then calculated Bray–Curtis distances (BCD) on the normalized k-mer counts using the ecodist R package (v. 2.1.3) (Goslee and Urban, 2007). We manually inspected genetic distances using principal coordinate analysis (PCoA) (Hrytsenko et al., 2025). For each dataset, we visually evaluated whether the main PCoA axes reflected the geographic patterns of genetic variation reported in the original studies (Figure S3E). In 3 out of 19 datasets (*Amphiprion bicinctus, Siphamia tubifer, Platygyra daedalea*), we observed strong genetic clustering along the first PCoA axis among samples from the same reef (*i*.*e*., sympatric clustering), and we therefore discarded samples from the smallest cluster (Figure S3, Table S1). Furthermore, we visually checked whether clusters of samples along the PCoA axes showed similar values for technical variables as mean read number, mean read length, GC content, and the percentage of duplicate reads. We found and discarded such clusters of samples in 5 out of 19 datasets (*Epinephelus striatus, Bartholomea annulata, Pocillopora verrucosa, Pinctada fucata* and *Carcharhinus amblyrhynchos*).

Finally, we assessed whether genetic distances based on k-mer counts were consistent with those based on traditional single nucleotide polymorphism (SNP)-based methods, specifically pairwise nucleotide diversity (π) ((Roberts et al., 2025; Roberts and Josephs, 2025). For this comparison, we focused on datasets from species with an available reference genome within the genus (16 out of 19 datasets), as reference genomes enable precise read alignment and reliable identification of SNPs. For each of these datasets, we randomly sampled 1,000 non-overlapping windows of 10,000 nucleotides from the reference genome using Bedtools (v. 2.31.1) ((Quinlan and Hall, 2010). Quality-filtered reads were then mapped to these reference genome segments using NextGenMap (v. 0.5.5) ((Sedlazeck et al., 2013). The mapped reads were sorted and indexed using Samtools (v. 1.16.1) ((Li et al., 2009), and both variant and invariant sites were called using Bcftools mpileup (v. 1.16) ((Li et al., 2009). We calculated π at each sampling location using the pixy method (v. 1.0.0) ((Korunes and Samuk, 2021), implemented in a custom R script (available in the online material) focusing on genomic windows with at least 100 genotyped nucleotides. To avoid bias in π estimates caused by uneven sample sizes across sites, we limited the analysis to five randomly selected samples per site. For each location, we also computed the average BCD among the same five samples based on k-mer counts and evaluated its correlation with π (Figure 2D).

**Figure 2.**
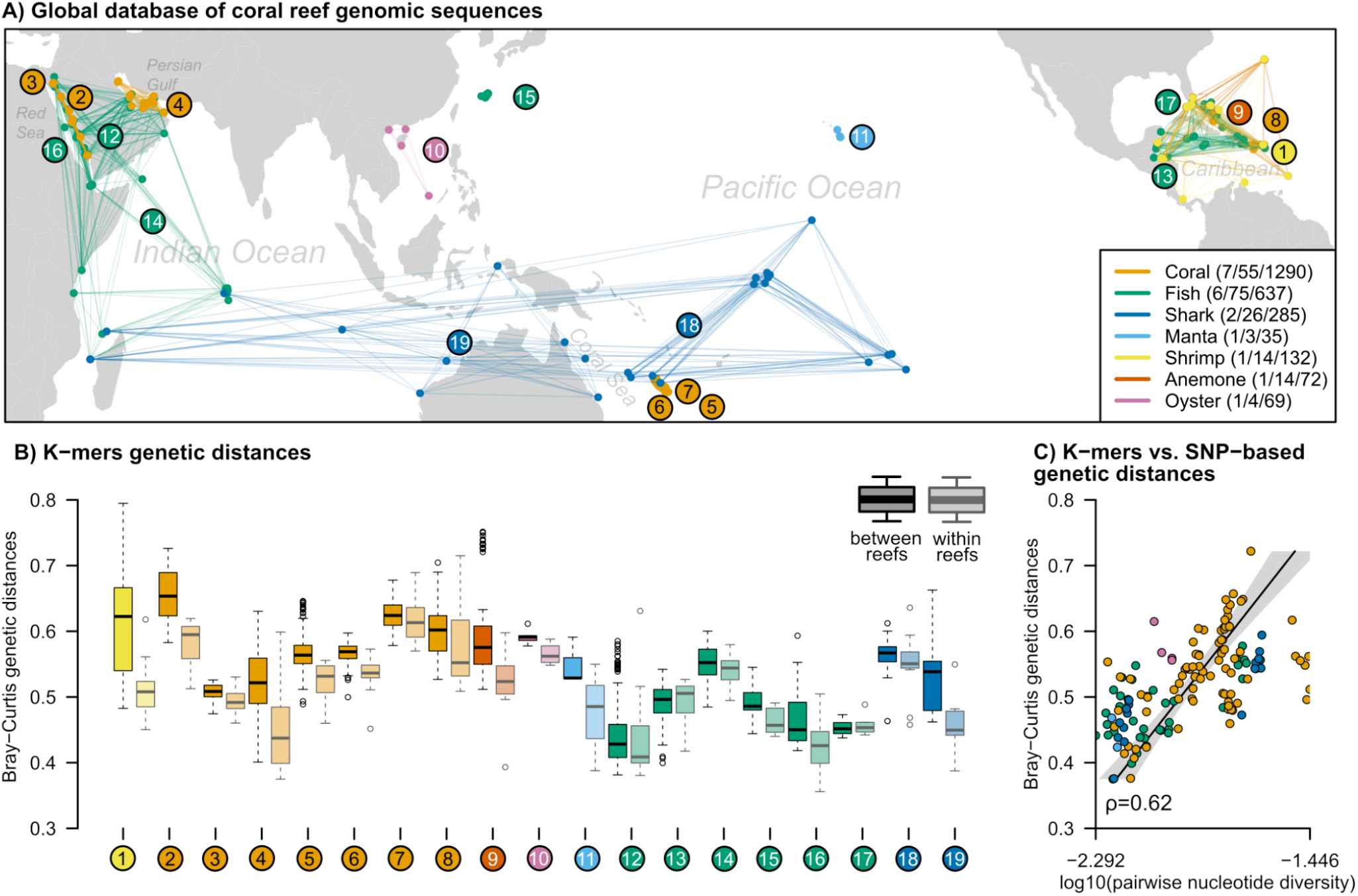
Genomic distances across the world’s reefs. (A) Geographic distribution of sampling locations across the 19 genomic datasets included in the study. Points represent sampling sites (N=173), linked by lines for each dataset (labeled from to 19, read below). The legend indicates the main taxonomic groups, with the number of datasets, sampling reefs, and individual samples shown in parentheses. For every dataset, the genetic distances between samples collected from different reefs (>10 km apart) and from the same reef (<10 km) are shown in (B). Genetic distances are calculated as Bray–Curtis distances (BCD) based on k-mer frequencies. (C) shows the correlation between within-reef BCD (calculated using k-mers) and pairwise nucleotide diversity (calculated from single nucleotide polymorphisms, SNPs). Correlation is reported using Spearman’s rank correlation coefficient (ρ). Datasets labels are: (1) *Ancylomenes pedersoni* from the Caribbean, (2) *Stylophora pistillata* and (3) *Pocillopora verrucosa* from the Red Sea, (4) *Platygyra daedalea* from the Persian Gulf, (5) *Pocillopora damicornis* (6) *Pocillopora acuta* and (7) *Acropora millepora* from New Caledonia, (8) *Acropora palmata* and (9) *Bartholomea annulata* from the Caribbean, (10) *Pinctada fucata* from the South China Sea, (11) *Mobula alfredi* from Hawaii, (12) *Pomacanthus maculosus* from the Indian Ocean, (13) *Pterois volitans* from the Caribbean, (14) Dascyllus trimaculatus from the Indian Ocean, (15) *Siphamia tubifer* from Japan, (16) *Amphiprion bicinctus* from the Red Sea, (17) *Epinephelus striatus* from Bahamas, (18 and 19) *Carcharhinus amblyrhynchos* from the Indo-Pacific.

### Characterization of coral reef environmental variation

The environmental characterization of the sampling locations was based on publicly available data compiled and standardized in the Reef Environment Centralized Information System (RECIFS, v. 2.0.1) ((Selmoni et al., 2023). RECIFS maps the global distribution of coral reefs (v. 4.1) ((Unep-Wcmc et al., 2021) using a 5 km grid, and for each grid cell provides longitude, latitude, reef area, and a set of environmental descriptors derived from satellite observations (complete list of product identifiers is available in Table S2). We matched the genomic sampling locations with the closest reef cell in RECIFS and extracted the associated environmental variables at two spatial scales: within the reef cell itself (2.5 km radius around the reef) and within the surrounding region (50 km radius).

Most of the environmental descriptors in RECIFS characterize water conditions, including thermal profile (sea surface temperature and degree heating weeks; from the NOAA Coral Reef Watch) (Liu et al., 2003; Skirving et al., 2020), chemical composition (chlorophyll, iron, nitrate, phosphate, oxygen concentration, sea surface salinity, and pH, from the Copernicus Marine Service) (EU Copernicus Marine Service, 2022), and physical properties (surface water velocity and suspended particulate matter, from the Copernicus Marine Service (EU Copernicus Marine Service, 2022). These variables are available at a monthly resolution over the past 2–3 decades, depending on the variable (see Table S2 for details). For each sampling location, we calculated yearly averages and minimum and maximum values of each variable. These time series were then summarized by computing the location’s overall mean, coefficient of variation, and rate of change (*i*.*e*., the difference between mean yearly values 1–10 years vs. 11–20 years before sampling).

In addition to time-resolved variables, we extracted RECIFS variables without monthly resolution that describe the surrounding seascape structure, such as average bathymetry and frequency of surface land (Ryan et al., 2009). We further complemented these data by calculating the reef area (in km^2^) surrounding each reef cell within buffer radii of 10, 50, 100, 250, and 500 km.

We also included RECIFS variables summarizing potential human impacts on reefs, including the frequency of built-up or cropland land cover (from the Copernicus Global Dynamic Land Cover dataset) (Copernicus Global Land Service, 2022), average human population density (CIESIN Columbia University, 2010), and average detection of boat traffic (Elvidge et al., 2018, 2015; Hsu et al., 2019). Finally, we retrieved the global distribution of Marine Protected Areas (MPAs) from the World Database on Protected Areas (WDPA; October 2023 release) ((UNEP-WCMC and IUCN, 2020) and calculated the distance from each reef cell to the nearest MPA.

### Modeling spatial and temporal patterns of genetic diversity

We used generalized linear mixed models (GLMMs) to characterize how coral reef genetic diversity varied across datasets, taxonomic groups, geographic regions, and periods of sampling. Models were built using the R package glmmTMB (v. 1.1.9) ((Brooks et al., 2017; Selmoni et al., 2023), applying a beta regression family to model the response variable: the BCD between each pair of samples within the same dataset (a total of 255,247 distance values, 51,259 of pairs from the same sampling location, 203,988 of pairs from different sampling locations).

Model construction followed a forward stepwise approach, using the Akaike Information Criterion (AIC) to evaluate model fit (Figure 3A, Table S3). A decrease of at least 2 AIC units was considered evidence of improved fit. Following this procedure, we progressively introduced variables into the model to explain variation in genetic distance between pairs of samples:

**Figure 3.**
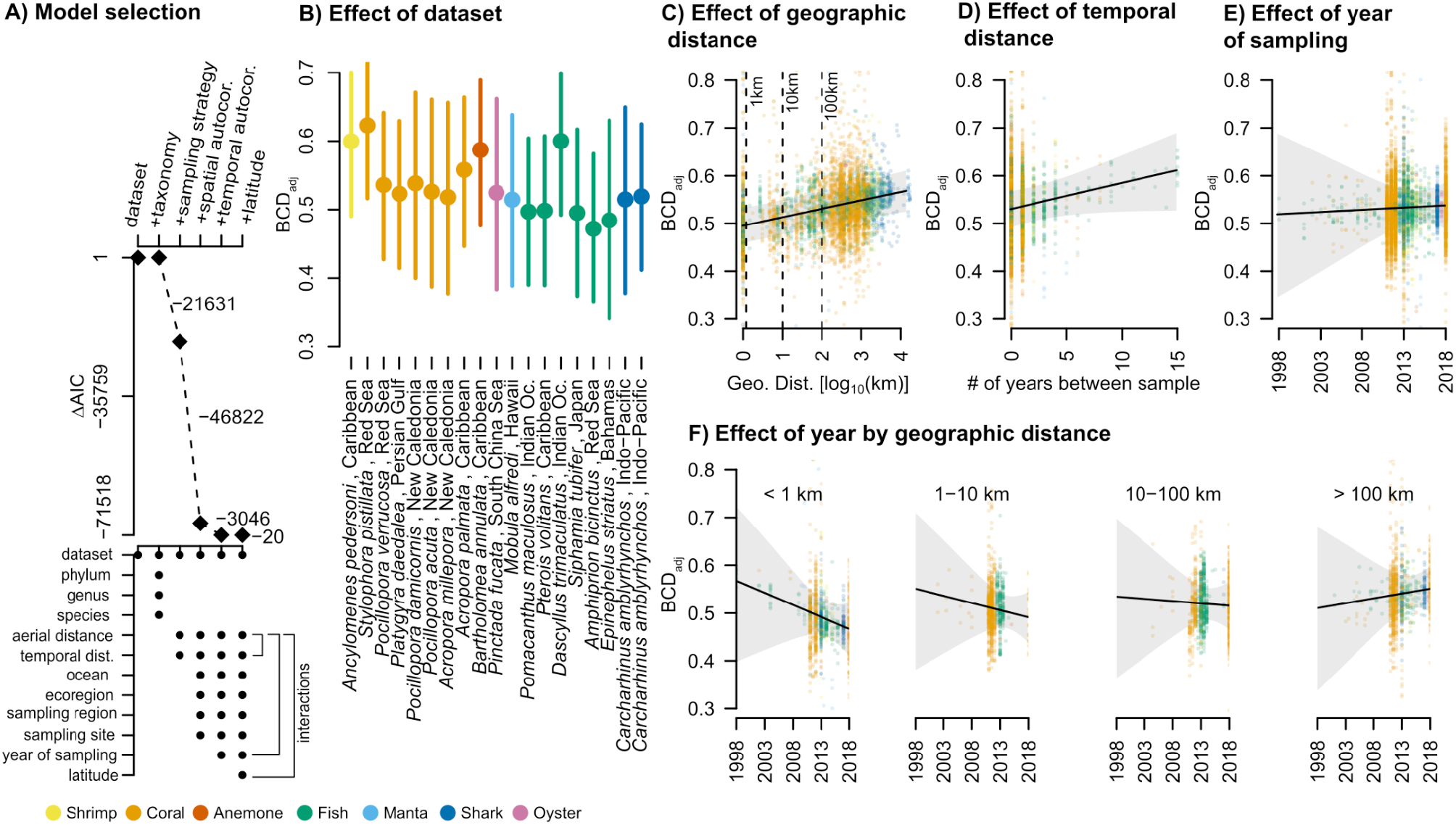
Taxonomic and spatiotemporal variation in coral reef genetic distances. (A) Stepwise model selection of variables explaining genetic distances (*i*.*e*. Bray-Curtis distances measured across 255,247 pairs of samples, from 18 coral reef species). Top: improvement in model fit (reduction of Akaike Inference Criterion, ΔAIC) relative to a null model including only dataset variation, for models incorporating taxonomy, sampling strategy, spatial autocorrelation, temporal autocorrelation, and latitude. Bottom names of variables included in each model. For the model with the best fit (lowest AIC, right side of A), (B-E) show the model-adjusted Bray–Curtis distances (BCD_adj_) for different variables, representing the marginal effect on genetic distances by dataset (B, points, with bars indicating the 95% confidence interval), geographic distance between sampling sites (C), temporal distance between years of sampling (D), and midpoint year of sampling (E). (F) BCD_adj_ by the midpoint year of sampling for sample pairs collected from different geographic distances (<1km, 1-10km, 10-100km, >100km). In (C-E), points represent the model’s partial residuals.

- Dataset: dataset identifier, random term (factor with 19 levels).
- Taxonomy: nested random terms for species, genera, and phyla (factors with 18, 15, 4 levels, respectively).
- Study design: number of years between sampling dates and aerial distance (log-transformed, in km) between individuals, fixed terms (continuous variables).
- Geographic autocorrelation: nested random terms for reef pairs (factor with 1440 levels, a reef groups samples located <10 km apart), region pairs (factor with 598 levels, individuals within 100 km), ecoregion pairs (factor with 136 levels, based on ecoregion polygons from the R package mregions2 (Fernández Bejarano and Pohl, 2024); v. 1.1.2), and ocean pairs (factor with 4 levels).
- Temporal autocorrelation: midpoint sampling year of the pair, fixed term (continuous variable).
- Latitude: absolute midpoint latitude of the pair, fixed term (continuous variable).

During stepwise model construction, we also tested two types of interactions. To account for potentially different patterns of genetic diversity within versus between populations, we included interaction terms between aerial distance and each of the following: years between sampling, year of sampling, and absolute latitude. Additionally, to capture dataset-specific variation, we allowed the random effect of the dataset to influence both the intercept and the slopes of aerial distance, years between sampling, and year of sampling.

The effects of the different variables on genetic distances were assessed using marginal effects calculated with the ggeffects (v. 1.3.4) ((Lüdecke, 2018) and emmeans (v. 1.10.2) ((Lenth and Piaskowski, 2025) R packages. For a given variable, the marginal effect represents the variable’s effect on model-adjusted BCD (BCD_adj_)–––*i*.*e* the predicted BCD while holding all other variables constant. The significance level of every fixed predictor was assessed with the built-in Wald test of glmmTMB. For every effect, we also evaluated significance by running a likelihood ratio test of the model with vs. without the effect of interest, using the anova function in the R car library (v. 3.1) (Fox et al., 2019) (Table S4).

### Matching local effects on genetic distances with environmental data

We investigated whether spatial patterns in coral reef genetic diversity could be explained by environmental variables derived from satellite observations. To do this, we used the GLMM defined in the previous section and extracted the local effect on genetic distances associated with every sampling site. For every sampling reef, this effect is the cumulative sum of the random variable effects of ocean, ecoregion, sampling region, and sampling reef (Figure S4). For example, the cumulative local effect for *Reef5*––located in sampling region *Region3*, ecoregion *Hawaii* and ocean *Pacific*–––is the sum of the random effects from pairs within *Reef5* pairs within *Region3* pairs within *Hawaii* and pairs within *Pacific*. Negative values of this cumulated effect indicate locations where genetic distances are lower than the global average, while positive values indicate locations where genetic distances are higher (Figure 4A).

**Figure 4.**
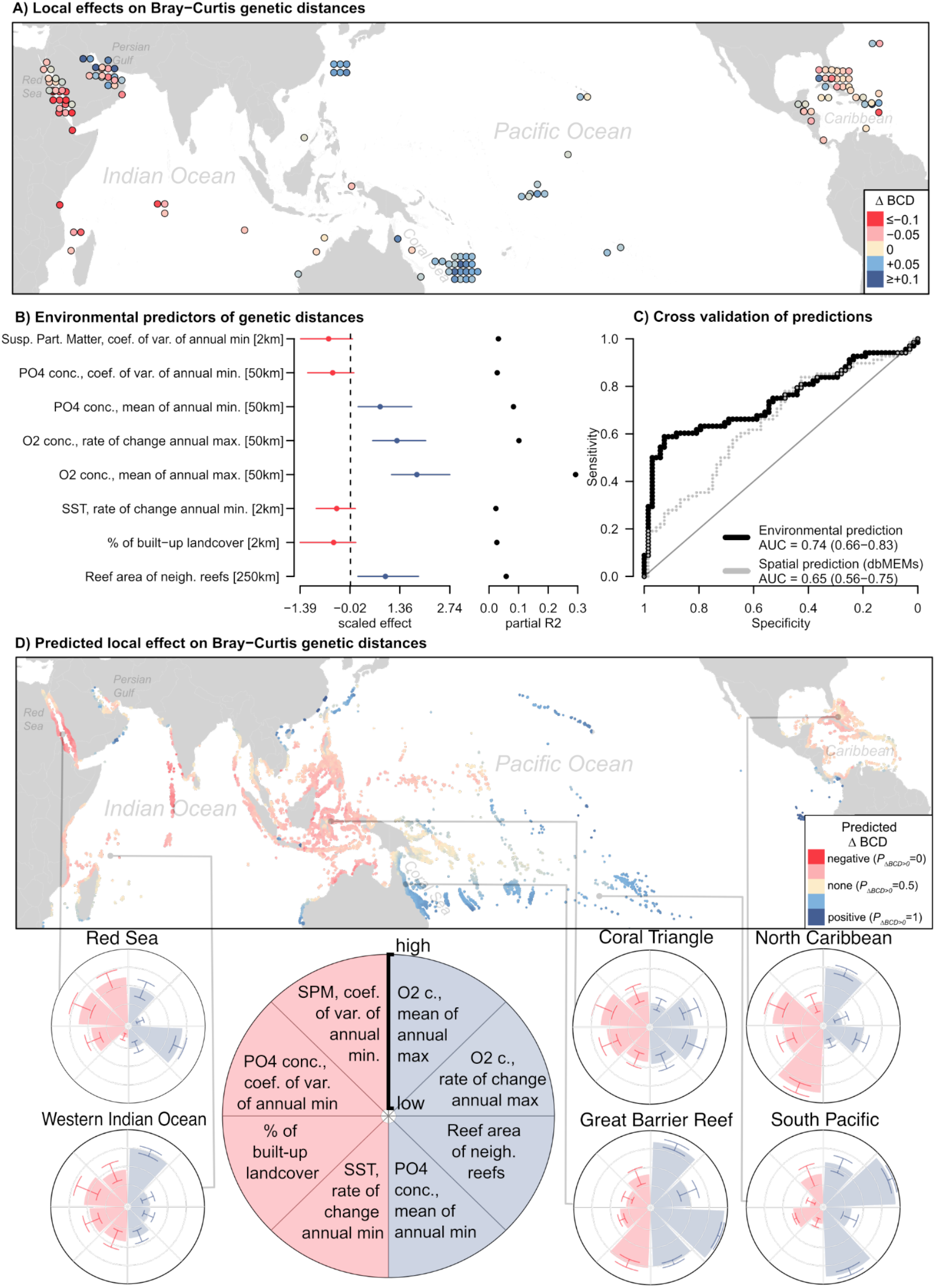
Prediction of genetic distances across the world’s coral reefs. (A) Cumulative local effects of geographic position o Bray–Curtis genetic distances at each sampling reef (N 136). Red indicates negative local effects o genetic distances; blue indicates positive effects. (B) Scaled effect sizes (with 95% confidence intervals) of the eight environmental variables retained in a penalized logistic model predicting local effects on genetic distances (in brackets: the spatial scale at which predictors were measured). Color reflects the direction of the effect (red negative, blue positive). Partia coefficients of determination (pR^2^) are shown on the right. (C) Predictive performance of the penalized logistic model under leave-one-region-out cross-validation, quantified using the Area Under the Curve (AUC). The black line shows the model based on environmental predictors; the grey line shows the performance of an analogous model that replaced environmental predictors with spatial predictors (distance-based Moran Eigenvector Maps, dbMEMs). (D) Global predictions of local effects o genetic distances cross coral reefs worldwide, based n the penalized logistic model Predicted probabilities of a positive effect (P_ΔBCD_ > 0) re shown: red ndicates likely positive effects (P_ΔBCD_ 1), blue negative effects (P_ΔBCD_ 0), and pale yellow indicates no predicted directional effect (P_ΔBCD_ 0.5). Bottom: Circular barplots display the scaled mean (± standard deviation) of the eight predictors shown in (B) across six focal reef regions. Bars are colored blue for variables associated with positive effects on genetic distances, and red for variables associated with negative effects.

We built a regression model using data from 136 sampling reefs across 19 datasets, with the local effect on genetic distance as response variable. Two alternative modeling approaches were tested:

1. A linear regression model, using the local effect on genetic distance as a continuous response variable.
2. A logistic regression model, using positive vs. negative local effects on genetic distance as a binary response variable.

For both models, we assessed the association between this response and 152 environmental variables characterized at the sampling reefs (as described in the “Characterize coral reef environmental variation” section). Given the large number of predictors, we used penalized regression (elastic net regularisation), implemented in the R package glmnet (v. 4.1-8) ((Friedman et al., 2010; Tay et al., 2023). This method applies penalties to predictor coefficients, shrinking the effects of less informative variables so that only the most relevant predictors are retained in the final model. The penalization is governed by two parameters: alpha, which controls how the penalty handles collinearity among predictors, and lambda, which determines the overall strength of penalization (Zou and Hastie, 2005).

We used cross-validation to identify the optimal values of alpha (tested across 10 levels from 0 to 1) and lambda (100 levels from 0.0001 to 0.2). For the linear model, the predictive performance was measured using the Mean Absolute Error between real vs. predicted effects on genetic distances (Figure S6). For the logistic model, we used the Area Under the Curve (AUC) metric, which evaluates whether the model discriminates correctly between positives vs. negative effects on genetic distance. Cross-validation was conducted using a leave-one-region-out strategy: in each iteration, the model was trained on sampling sites from all but one region and then used to predict local effects in the left-out region. Regions were defined by grouping sampling sites located within 1,500 km, which approximates the average size of Large Marine Ecosystems (estimated using the R package mregions2, v. 1.1.2)) (Fernández Bejarano and Pohl, 2024). To assess how predictive performance changes with spatial scale, we also tested two alternative region sizes: ecoregion scale (<800 km) and marine realm scale (<5,000 km; Figure S5).

To evaluate the contribution of spatial autocorrelation to model performance, we built an alternative version of both the linear and the logistic models, in which environmental variables were replaced with Moran Eigenvector Maps (MEMs)—orthogonal variables describing spatial proximity between sampling sites. These were calculated using the adespatial R package (v. 0.3-23) ((Dray et al., 2025). The same penalized regression and cross-validation procedure was applied to test whether geographic position alone (without environmental information) could predict local effects on genetic distances (Figure S7). Results showed that the linear model performed similarly with environmental and spatial variables (Figure S6), unlike the logistic model (Figure 4C). Consequently, we prioritized the logistic model for further analysis of explanatory variables.

We characterized the association between top environmental predictors and positive vs. negative effects on genetic distances. Because some of the predictors retained by the regularized regression may be redundant, we removed those that did not improve model fit using backward stepwise model selection based on the Akaike Information Criterion (AIC) (via the step function in the stats package, v. 4.3.2). The effects of the retained environmental variables were then visualized using the visreg R package (v. 2.7.0) ((Breheny and Burchett, 2017). In addition, we evaluate the percentage of variance explained by each of these environmental variables using the partial coefficient of determination, calculated using the rsq R package (v. 2.7) ((Zhang, 2024).

### Predicting local effects on genetic distances for reefs worldwide

To map local effects on genetic distances across the world’s coral reefs, we used environmental predictors to estimate the probability of a positive local effect on genetic distances (P_ΔBCD_ > 0). Specifically, we retrieved the set of predictors retained by the penalized logistic model and extracted their values across all global reef cells from RECIFS (Selmoni et al., 2023), following the same procedures described in the “Characterize coral reef environmental variation” section. We then applied the trained model to generate predictions of P_ΔBCD_ for each reef cell. Predicted values close to 1 indicate reefs likely to exhibit positive local effects on genetic distances, values near 0 indicate likely negative effects, and values around 0.5 suggest no clear directional trend. To help interpret geographic variation in these predictions, we summarized regional patterns of environmental drivers using circular barplots generated with the ggplot R package (v. 3.5.1) ((Wickham, 2016), highlighting how key seascape conditions differ across reef regions.

## 3. Results

### Kmer-based genetic distances across the world coral reefs

We built a comprehensive database to compare genetic diversity in coral reef species from different regions of the world (Figure 2A). The database included 19 datasets spanning a variety of taxa, including corals (seven datasets), fish (six), sharks (two), manta ray, shrimp, oyster, and sea anemone (one each). In terms of geography, five datasets covered reefs in the Atlantic Ocean, six in the Pacific Ocean, six in the Indian Ocean, and two in the Indo-Pacific. Each dataset covered a median spatial scale of ~3,000 km (interquartile range, IQR ≈ 2,600 km). Seven datasets were sampled in a single year while the others spanned two or more years, with sampling years, ranging from 1998 to 2018 (Figure S8). The initial sample size was 3,037 individuals (range per dataset: 37 to 448), of which 2,520 (range: 28 to 393) were retained after quality control, covering 173 coral reefs globally (Table 1, Table S1).

In every dataset, we assessed genetic distances between pairs of samples applying the Bray–Curtis method to k-mers frequencies (Roberts et al., 2025) (Figure 2B). Mean Bray-Curtis genetic distances (BCD) ranged from 0.44 to 0.63 across datasets, and were generally lower between pairs from the same reef (sampling sites <10 km apart) compared to pairs from different reefs (>10 km apart; difference in mean BCD ranging −0.11 to 0.01; Figure 2B, Table S1).

To ensure that k-mer-based genetic distances reflected true biological patterns, we applied two plausibility checks. First, we inspected the structure of the BCD matrices using Principal Coordinate Analysis (Hrytsenko et al., 2025) (PCoA, Figure S3), which confirmed that observed population structures aligned with geographical separation between sampling sites and were not driven by technical artifacts (*e*.*g*., sequencing reads abundance and quality) or sampling errors (*e*.*g*., cryptic species in sympatry). Second, for datasets with an available reference genome, we calculated pairwise nucleotide diversity (π) based on SNPs and found a significant correlation between π and k-mer-based distances within reefs (Spearman’s rank correlation ρ=0.62, *P<2*.*2e-16*; Figure 2B).

### Disentangling spatiotemporal patterns of coral reef genetic diversity

We aimed to characterize how genetic diversity across the world’s coral reefs varies with taxonomic group, geographic region, and year of sampling, while accounting for variation among datasets and sampling strategies (Figure S9). To do so, we built a generalized linear mixed model (GLMM; Figure 3) describing BCD between all pairs of samples within each dataset for a total of 255,247 pairwise comparisons across all datasets. Starting from a null model that accounted only for dataset-level variation, we used forward stepwise selection to identify explanatory variables that improved model fit as evaluated using the Akaike Information Criterion (AIC; Figure 3A; Table S3). The final model retained variables characterizing:

- Sampling strategy: geographic distance between sampling sites and temporal separation between sampling years of each pair of samples (ΔAIC = −21,361 compared to the null model);
- Geographic autocorrelation: identifiers for oceans, ecoregions, sampling regions, and sampling reefs of each pair (ΔAIC = −46,822);
- Temporal autocorrelation: midpoint year of sampling for each pair (ΔAIC = −3,046);
- Latitudinal effects: absolute midpoint latitude of each pair (ΔAIC = −36).

Taxonomy (species, genus, phylum) did not improve model fit (ΔAIC ≈ 0), likely because taxonomic variation was already captured by differences between datasets.

To interpret how each retained variable contributed to genetic distance variation, we calculated model-adjusted BCD (BCD_adj_). BCD_adj_ represents the marginal effect of a focal explanatory variable on genetic distance, while holding all other variables constant (Figure 3B–F; Tables S2–S3).

#### Effects of datasets and sampling strategy

Genetic distances varied significantly between datasets (χ^2^[23] = 852980 *P* < 2.2 10 ^16^), with invertebrates showing a higher median BCD_adj_ of 0.54 (IQR = 0.05), compared to vertebrates (0.50 IQR = 0.02; Figure 3B). Genetic distance increased significantly with geographical distance between sampling sites. Specifically, BCD_adj_ increased by 0.018 per log_10_(km) of separation, *i*.*e*. on a linear-log scale (95% confidence interval, CI: 0.01 to 0.025; *P* = 5.97 × 10^-6^ Figure 3C). This trend varied across taxa: invertebrates showed twice the rate of increase with distance compared to vertebrates (median slope = +0.02 BCD_adj_ per log_10_-(km) [IQR = 0.02] for invertebrates *vs*. +0.01 [IQR = 0.01] for vertebrates; Figure S10A).

We also detected a weak overall association between genetic distance and temporal separation between sampling years: across all sample pairs, BCD_adj_ increased by +0.0055 per year (95% CI: –0.0004 to 0.011, *P* = 0.08; Figure 3D). However, the effect of temporal separation varied between pairs collected across different spatial scales (interaction *P* = 8.08 × 10^−23^ Figure S11). In same-site pairs (<1 km apart), genetic distances increased strongly with temporal distance (+0.014 BCD_adj_ per year [95% CI: 0.008 to 0.02], *P* = 7.83 × 10^−6^), while this association became progressively weaker in pairs collected over greater distances (*e*.*g*, for samples > 100 km apart: +0.004 BCD_adj_ per year [95% CI: –0.002 to 0.009], *P* = 0.19; Figure S11; Table S5A).

#### Genetic distance change over time

We did not observe a significant overall trend in genetic distance change over time when considering all sample pairs within each dataset (Figure 4E). However, trends varied across datasets although this variation was not clearly attributable to taxonomy (range of median BCD_adj_ in invertebrates: –0.05 to 0.01, in vertebrates –0.003 to 0.03; Figure S10C). We then observed strongly contrasting trends depending on the spatial scale of the sample pairs (interaction *P* = 7.71 × 10^−10^ Figure 4F). For example, within-site sample pairs (<1 km apart) showed a negative trend in genetic distance over time (–0.0051 BCD_adj_ per year [95% CI: –0.016 to 0.0054]), while pairs separated by more than 100 km exhibited a slight positive trend (+0.002 BCD_adj_ per year [95% CI: –0.0084 to 0.012]; Table S5B).

#### Geographic patterns of genetic distances

Genetic distances showed a weak association with latitude (–0.0018 BCD_adj_ per degree of latitude [95% CI: –0.0036 to 0], *P* = 0.04; Figure S12, Table S5C), but varied significantly across all geographic scales: oceans (χ^2^[23] = 7, *P* = 0.006), ecoregions (χ^2^[23] = 109, *P* < 2.2 × 10^−16^), sampling regions (χ^2^[23] = 19764, *P* < 2.2 × 10^−16^), and sampling sites (χ^2^[23] = 62, *P* = 2.9 × 10^−15^ Figure S4; Table S4). On average, local effects on genetic distances were negative for sample pairs from the Indian Ocean (ΔBCD= –0.03) and positive for those from the Atlantic Ocean (ΔBCD= +0.01) and the Pacific Ocean (ΔBCD = +0.02; Figure S4A).

When focusing on each of the finer spatial scales, geographic patterns were more difficult to generalize, although negative effects were more frequently observed in pairs from the Northern Indian Ocean and the Caribbean (Figure S4B–D, Table S6). However, spatial patterns became more evident when we aggregated spatial effects across all levels—ocean, ecoregion, sampling region, and sampling reef—and summed such effects for each sampling site (Figure 4A). This revealed consistently negative effects in sampling sites from the Red Sea and Persian Gulf (median ΔBCD = −0.03 [IQR = 0.08]) and the Northern Caribbean (median ΔBCD = –0.02 [IQR = 0.03]), and positive effects in the Coral Sea (median ΔBCD = +0.05 [IQR = 0.03]).

### Predicting coral reef genetic diversity from seascape conditions

We finally asked whether the genetic diversity of a reef in unsampled regions can be predicted from its seascape conditions. For this, we extracted local effects on genetic distance across 136 sampling reefs (Figure 4A), and then built a penalized logistic model using 152 seascape variables to discriminate between reefs with positive or negative local effects.

#### Model predictive performance

The model correctly discriminated positive vs. negative effects on genetic distances 74% of the time (Area Under the Curve [AUC]= of 0.74 [95% CI: 0.66 to 0.83]; Figure 4C). This result was obtained using a leave-one-region-out cross-validation, where regions were defined at the scale of Large Marine Ecosystems (reefs located up to 1,500 km apart; Figure S5B). We then repeated the cross-validation at the finer scale of ecoregions (~800 km), obtaining a minor improvement in performance (AUC = 0.77 [95% CI: 0.68 to 0.85]; Figure S5A). We also tested the model’s ability to extrapolate to larger unsampled regions, and repeated the cross validation at the marine realm scale (~5,000 km). In this case, the model performed no better than random (AUC = 0.51 [95% CI: 0.41 to 0.61]; Figure S5C).

To verify whether the model’s predictive power was driven by spatial autocorrelation, we built an alternative model using only Moran Eigenvector Maps (MEMs) ((Dray et al., 2006) as predictors. The performance of this model was better than random but lower than our environmental model (AUC = 0.65 [95% CI: 0.58 to 0.75]; Figure 4C; Figure S7).

#### Seascape conditions explain the geographic variation of genetic distances

We then wanted to identify which seascape conditions best explain geographic patterns of genetic diversity. The penalized logistic model retained eighteen seascape variables that optimized predictions of genetic distances (Figure S13). These variables captured a range of seascape characteristics, including oxygen and phosphate concentrations, sea surface temperature, suspended particulate matter, reef area, and proximity to built-up landcover.

We further restricted this list using stepwise regression and identified eight non-correlated predictors that best explained geographic variation in genetic distance (model R^2^ = 0.52; Figure 5B; Table S7; Figure S14). Two selected predictors explained a substantial portion of this variation, both related to annual maximum oxygen concentration: negative effects on genetic distances were observed at sites with low oxygen concentration (partial R^2^ pR^2^ = 0.29), and at sites where oxygen concentration had decreased over recent decades (pR^2^ = 0.10). Other retained predictors explained smaller fractions of the variation. Two were related to annual minimum phosphate concentration: high concentrations were associated with locally increased genetic distances (pR^2^ = 0.08), while higher interannual variability was linked to decreased genetic distances (pR^2^ = 0.03). The surface area of neighboring reefs (*i*.*e*, within 250 km of a focal reef) also contributed to explaining geographic variation in genetic distances (pR^2^ = 0.06), with larger reef areas associated with positive local effects. Another retained predictor was the annual minimum of suspended particulate matter, with decadal variation showing a negative association with genetic distances (pR^2^ = 0.03). The proportion of built-up landcover within 2 km of a reef was also associated with negative local effects on genetic distances (pR^2^ = 0.03). Last, reefs where the annual minimum SST increased over recent decades tended to exhibit negative effects on genetic distances, but the geographic variance explained by this SST variable was relatively low (pR^2^ = 0.02).

#### A global predictive map of coral reef genetic diversity

To assess how seascape conditions may influence genetic diversity globally, we used our model to predict local effects on genetic distances for every coral reef worldwide (Figure 4D). The predicted value was the probability of a positive local effect on genetic distance (P_ΔBCD_ > 0), where values close to 1 indicate a predicted positive local effect on genetic distances, values near 0 indicate a negative effect, and values around 0.5 indicate no expected directional effect.

At the ocean basin scale, predicted negative effects on genetic diversity were most frequent in the Indian Ocean (median P_ΔBCD_ > 0 = 0.33 [IQR = 0.17]), whereas predictions for the Atlantic (0.41 [IQR = 0.11]) and Pacific (0.48 [IQR = 0.37]) showed no clear directional pattern. However, distinct subregional trends emerged within each ocean. For instance, the Red Sea (P_ΔBCD>0_ = 0.23 [IQR = 0.14]), the North Caribbean (0.37 [IQR = 0.09]) and the Coral Triangle (P_ΔBCD>0_ = 0.32 [IQR = 0.09]) were predicted to have predominantly negative effects on genetic distances. In contrast, positive effects on genetic distance were predicted for the Great Barrier Reef (P_ΔBCD>0_ = 0.74 [IQR = 0.13]) and South Pacific Islands (0.77 [IQR = 0.05]).

## 4. Discussion

### K-mers distances as a tool for macrogenomics

The results of this study show that k-mers are a promising tool for macrogenetic studies. Our computational pipeline characterized genetic distances across hundreds of samples from diverse species. The only deviation from the standard pipeline was the k-mer filtering strategy for species without a reference genome. However, the alternative approach (discarding k-mers matching contaminant genomes) did not yield systematic differences in observed genetic diversity patterns. Our analysis confirms the relationship between BCD and π recently reported in plants (Roberts and Josephs, 2025), supporting k-mer-based distances as a practical and computationally efficient approach for assessing genetic diversity without mapping entire DNA sequencing libraries against a reference genome (Hrytsenko et al., 2025).

K-mer-based genetic distances were consistent with known taxonomic differences. For example, genetic distances were larger in invertebrate datasets, aligning with previous research showing that marine invertebrates—such as corals, anemones, shrimp, and oysters—generally exhibit stronger genetic diversity than vertebrates, such as fish and sharks (Toczydlowski et al., 2025). Furthermore, the increase in genetic distance with geographic separation was twice as strong in invertebrates as in vertebrates. This may reflect differences in dispersal potential: vertebrates often benefit from both adult movement and gamete dispersal, whereas invertebrates typically rely on gamete dispersal alone (Hernawan et al., 2021; Selkoe et al., 2014).

### Are coral reef populations rapidly losing their genetic diversity?

Despite widespread coral reef degradation over recent decades (Souter et al., 2021), we did not detect a clear overall trend in genetic diversity changes over time. Instead, trends varied across datasets without a clear taxonomic link. This aligns with recent evolutionary simulations showing that, shortly after habitat loss, species-wide genetic diversity can either increase or decrease, depending on the geographic pattern of habitat loss (Mualim et al., 2026).

The same simulations predicted that temporal trends in genetic diversity differ between population (deme) and species levels (Mualim et al., 2026). Consistently, we observed contrasting temporal trends depending on spatial scale: genetic diversity declined in pairs from the same reef during 1998-2018, while it showed a slight increase in distant pairs (sampled >100 km apart). This could be explained by the Wahlund effect, where reduced gene flow between reefs increases genetic differentiation among populations but reduces within-reef diversity.

Previous studies have found little evidence of genetic diversity declines in coral reefs over recent decades (Pinsky et al., 2023). Notably, while our within-reef analysis suggests a relatively strong negative trend (approximately 18% decline in BCD_adj_ over 20 years), this trend remains uncertain and not statistically significant. Additionally, we could not assess how this trend varied across taxa and regions. This uncertainty likely stems from limited temporal coverage: only half of our datasets include multi-year samples, and very few predate 2010. Future work could address this by incorporating more recent genetic data (post-2018) to test whether the observed negative trend is confirmed across regions and taxa.

### Where and how climate shapes coral reef genetic diversity

The spatial distribution of genomic studies on coral reefs remains highly unbalanced (Pinsky et al., 2023). Our database reflects this, with dense sampling in the Red Sea, Persian Gulf, Coral Sea, Japan, and the Northern Caribbean, while vast regions—such as Southeast Asia, Pacific Small Island States, and the Southern Caribbean—are largely unsampled (Figure 2A). Our study demonstrates that satellite-derived seascape variables can help predict genetic diversity patterns in unsampled areas, with some limitations. First, our model could distinguish reefs with positive vs. negative local effects on genetic diversity but could not quantify their magnitude. Second, predictions were reliable only when training data existed within the same Large Marine Ecosystem (~1,500 km from the focal site). Extrapolating the model across larger biogeographic breaks (*e*.*g*., into new marine Realms) yielded random predictions. Together, these findings highlight the potential of environmental data to interpret and predict global genetic diversity patterns better than geographic proximity alone, while suggesting that predictive power could improve with more globally balanced training data.

The predicted negative effects on genetic diversity in the Red Sea and North Caribbean are particularly notable, as both regions are well-represented in our dataset with multiple taxa sampled across numerous reefs. In both areas, declining oxygen concentrations over recent decades emerge as a shared explanatory factor. Hypoxia is a recognized threat to marine biodiversity and a growing concern for coral reefs (Hughes et al., 2020; Johnson et al., 2021). A recent global study found that one-third of surveyed reefs are already exposed to moderate hypoxia, with projections indicating this could rise to over half by 2100 (Pezner et al., 2023). Hypoxia tolerance varies among coral reef species and can drive shifts in community diversity (Hughes et al., 2020; Johnson et al., 2021; Lucey et al., 2024). In the North Caribbean, negative local effects on genetic diversity may also be linked to rising annual minimum sea surface temperatures, as marine heatwaves are a major driver of coral reef degradation (Hughes et al., 2017). The Coral Triangle is another region predicted to experience negative local effects on genetic diversity, although these predictions should be interpreted cautiously due to the lack of training data from this region. These negative predictions likely result from persistently low oxygen concentrations, high coastal built-up land cover density, and variable phosphate levels. While phosphate is essential for coral reef health (Ezzat et al., 2016), anomalously high or fluctuating levels can lead to eutrophication and reef degradation (Silbiger et al., 2018).

We also identified regions with predicted positive effects on genetic distance, like the Great Barrier Reef and South Pacific Islands. In the Great Barrier Reef, the negative impact of increasing minimal annual temperatures may have been offset by persistently high oxygen levels and large reef areas. This aligns with theoretical expectations that larger habitats maintain higher genetic diversity (Exposito-Alonso et al., 2022) and empirical evidence from Hawaii showing a positive correlation between coral reef area and species-level genetic diversity (Selkoe et al., 2014). In the South Pacific, stable oxygen and temperature regimes over recent decades likely support the observed positive predictions.

Previous macrogenetic studies on terrestrial species have revealed a clear latitudinal gradient in genetic diversity, with diversity declining toward the poles (Miraldo et al., 2016; Yiming et al., 2021). In marine species, similar trends have been observed in studies based on mitochondrial markers (Manel et al., 2020), but not when using nuclear DNA (Clark and Pinsky, 2024; Figuerola-Ferrando et al., 2023). Our results suggest that coral reef genetic diversity might follow this latitudinal gradient, although the signal is weaker than in the previous works. Furtermore, our predicted negative spatial patterns of genetic diversity differ from those reported in previous oceanic macrogenetic studies (Clark and Pinsky, 2024; Figuerola-Ferrando et al., 2023; Manel et al., 2020). For instance, while the Coral Triangle has been identified as a genetic diversity hotspot for fish (Clark and Pinsky, 2024; Manel et al., 2020), and the North Caribbean for habitat-forming taxa (Figuerola-Ferrando et al., 2023), our model predicted negative local effects in both regions. These discrepancies likely stem from methodological differences: our study is the first to focus specifically on coral reef species and to use genome-wide DNA sequencing data, rather than nuclear microsatellites or mitochondrial markers. Marker choice is a known source of variation in macrogenetic studies (Paz-Vinas et al., 2021), and further comparisons are needed to assess consistency across marker types.

Taken together, these results can help prioritize coral reef areas for field sampling and DNA-based genetic diversity monitoring. For instance, Earth observation datasets can track near-real-time changes in oxygen and phosphate concentrations across global reefs. Oxygen levels or phosphate concentrations can be used as informative factors to design sampling strategies. Sampling should furthermore account for reef surface area, as smaller surface areas are associated with lower genetic diversity.

## 5. Notes

## Acknowledgements

We are grateful to the openness of many researchers who make genomic data publicly available and this research possible: Buitrago-Lopez et al., Smith et al., Selmoni et al., Devlin-Durante et al., Titus et al.,Shan et al., Whitney et al., Torquato et al., Bors et al., Salas et al., Gould and Dunlap, Saenz-Agudelo et al., Sherman et al., Lesturgie et al., Walsh et al.. We thank members of the Spatial Genetics Group for comments and discussions. We furthermore thank M. E. Schaepman for supporting this work, B. Schmid for the suggestions on statistical modeling, and the Spatial Genetics of Ecosystem laboratory for the helpful discussions.

## Author contributions

OS: Conceptualization, data curation, formal analysis, funding acquisition, investigation, methodology, visualization, writing (original draft). MS: resources, supervision writing (review and editing).

## Data availability

The analyzed datasets are publicly available (see Supplemental Table 1 for data links). Processed data are available at Zenodo (doi: 10.5281/zenodo.19205727) and Github (https://github.com/Oselmoni/GFS_coralreefs).

## Code availability

Code to reproduce the analysis is available at Zenodo (doi: 10.5281/zenodo.19205727) and Github (https://github.com/Oselmoni/GFS_coralreefs).

