## Supplementary Figures for "Spatiotemporal patterns of genetic diversity in the world’s coral reefs"

This document includes Supplementary Figures 1-13.

**Table of content:**

Supplementary Figure 1. Distribution of sequencing depth across datasets. 2
Supplementary Figure 2. Filtering statistics and population genetic structure of the datasets. 3
Supplementary Figure 3. Geographic effects on genetic distances. 23
Supplementary Figure 4. Leave-one-region-out cross-validation of logistic elastic net regularization models. 24
Supplementary Figure 5. Leave-one-region-out cross-validation of linear elastic net regularization models. 25
Supplementary Figure 6. Elastic net regularization model using spatial predictors. 26
Supplementary Figure 7. Spatial and temporal distribution of datasets. 27
Supplementary Figure 8. Unadjusted associations between genetic distances and sampling design variables. 28
Supplementary Figure 9. Dataset-specific effects on genetic distances. 28
Supplementary Figure 10. Effect of the interaction between aerial and temporal distance on genetic distances. 30
Supplementary Figure 11. Effect of the interaction between aerial distance and latitude on genetic distances. 31
Supplementary Figure 12. Environmental predictors retained after elastic net regularization. 32
Supplementary Figure 13. Environmental predictors associated with local effects on genetic distances. 33
Supplementary Figure 14. Checking the effect of downsampling DNA sequencing reads. 34

**Supplementary Figure 1. Distribution of sequencing depth across datasets.**

For each dataset, the plot shows the distribution of the number of reads per sample (in millions). Red bars indicate the 10th percentile of the distribution, representing the threshold used for downsampling reads.

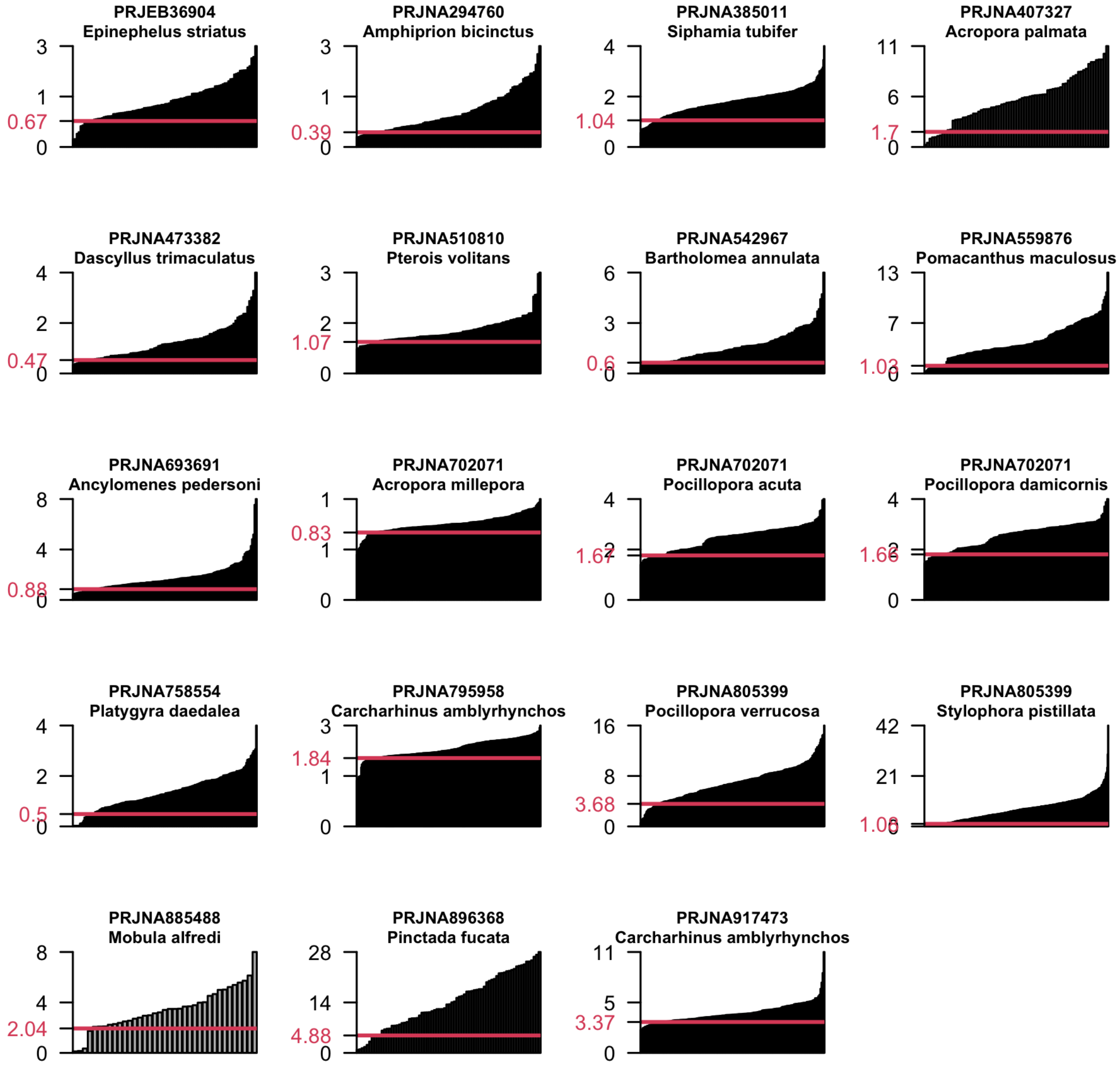

**Supplementary Figure 2. Checking the effect of downsampling DNA sequencing reads.**

Shown are the results of Principal Coordinate Analyses (PCoA) based on k-mer counts for a grey reef shark dataset. In (A), k-mers were counted on all DNA sequencing reads. In (B), reads were downsampled to the 10th percentile of DNA library size distribution. The maps show the geographic distribution of samples, colored by their position on the PCoA axes (shown in the inset, with the percentage of variance explained by each axis).

**A - Population structure using all reads**

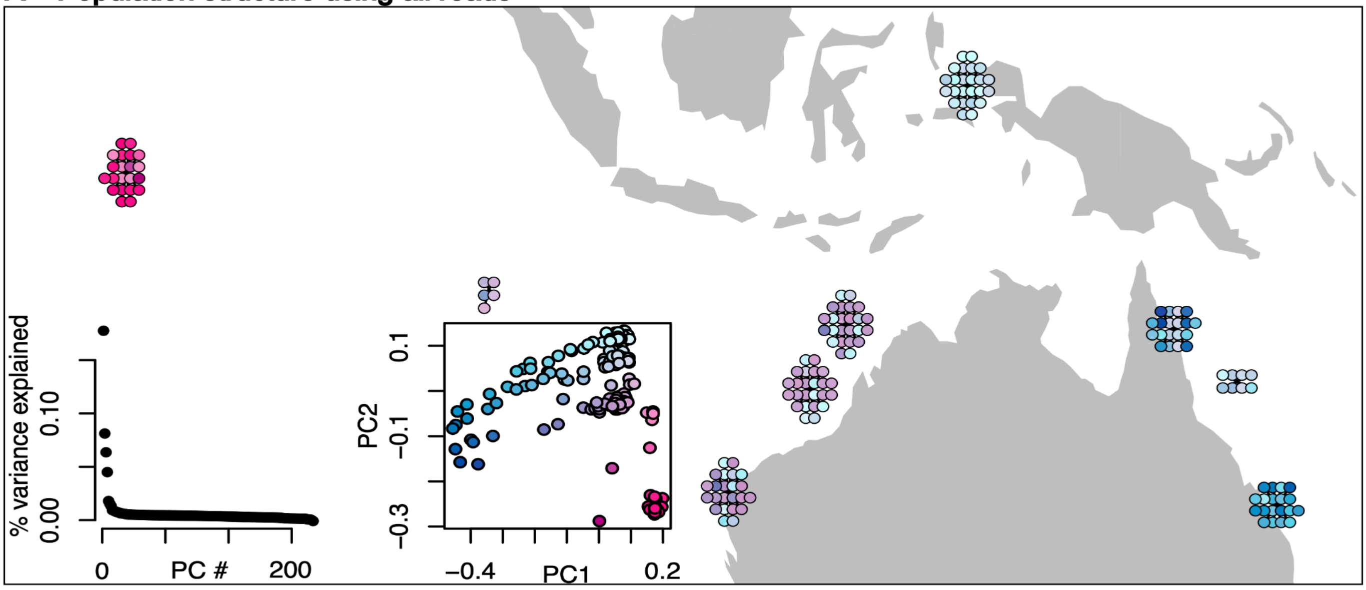


**B - Population structure using the 10th percentile of reads distribution**

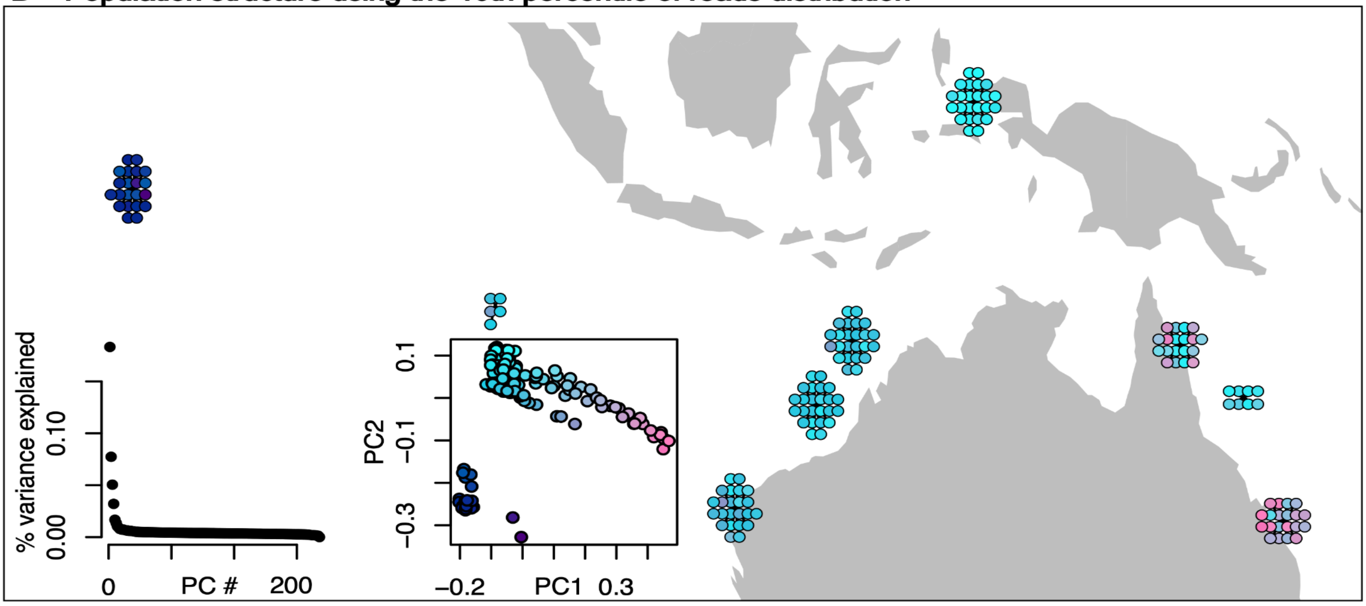

**Supplementary Figure 3. Filtering statistics and population genetic structure of the datasets.**

For each of the 19 datasets (each displayed on a different page), multiple panels summarize quality filtering outcomes and population structure. The name of each dataset (with BioProject ID and species name) is shown at the top of each page. For each dataset, the map displays the spatial distribution of genotyped individuals retained after quality filtering, colored by the first two axes of the Principal Coordinate Analysis (PCoA) of the k-mer matrix; symbol shapes represent sampling years. Panels show the following: (A) GC content of sequencing reads retained vs. excluded after taxonomic filtering; (B) Sample-wise GC content distribution, with red areas indicating outlier samples that were discarded; (C) Frequency distribution of k-mers, with red areas marking reads excluded for falling outside the 5th–95th percentile thresholds; (D) Correlation of k-mer frequencies between samples, red areas indicate highly similar samples (Spearman's $\rho > 0.9$), flagged as potential clones; (E) Preliminary PCoA used to detect technical artifacts (e.g., structure associated with sequencing statistics or cryptic species); samples in red were excluded; (F) Final PCoA showing the percentage of variance explained and the distribution of samples along the first four axes; (G) Spearman correlations between sequencing statistics (read depth, duplication rate, GC content, and read length) and the first two PCoA axes.

**Figure 2 (continue)**

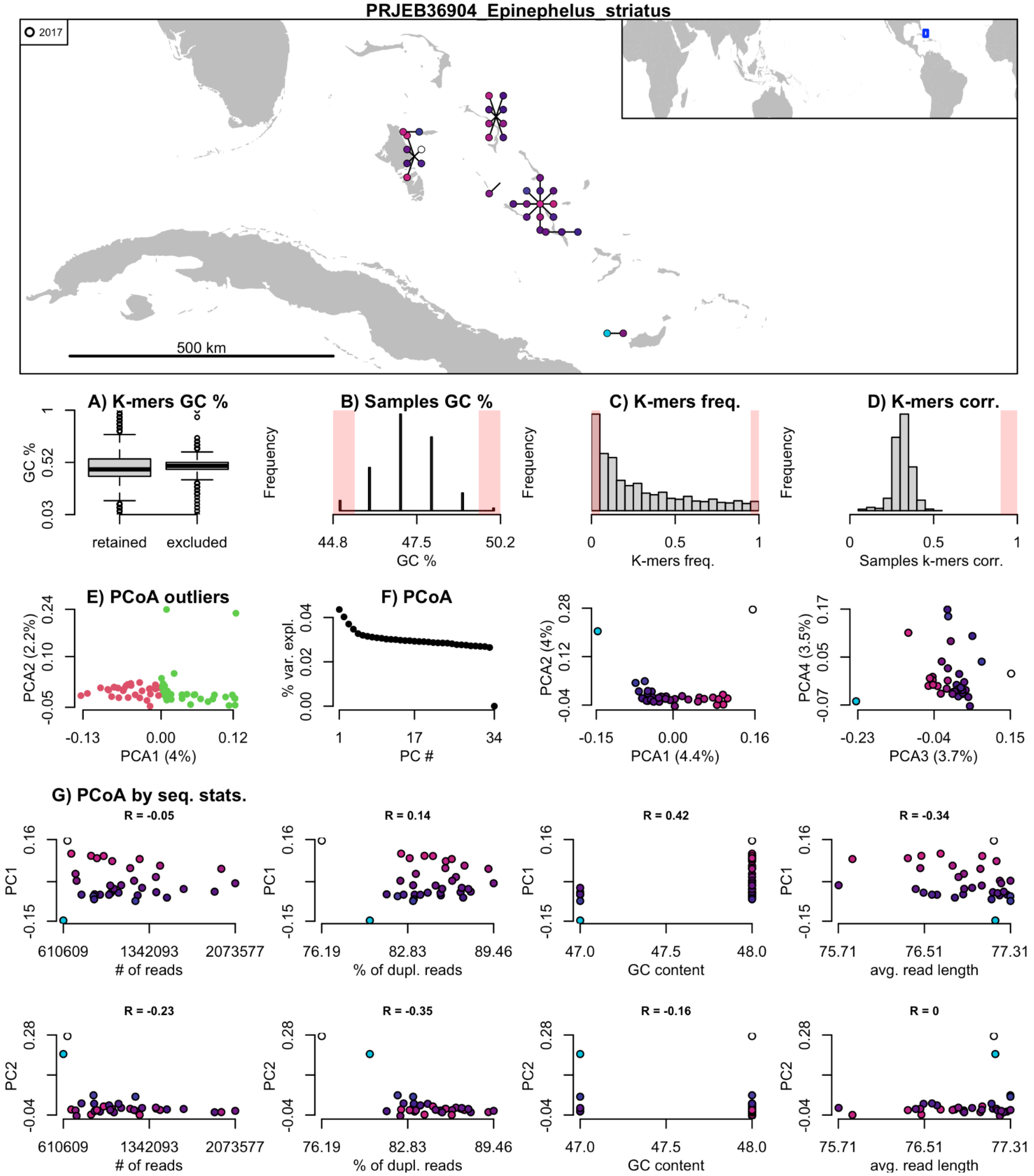

**Figure 2 (continue)**

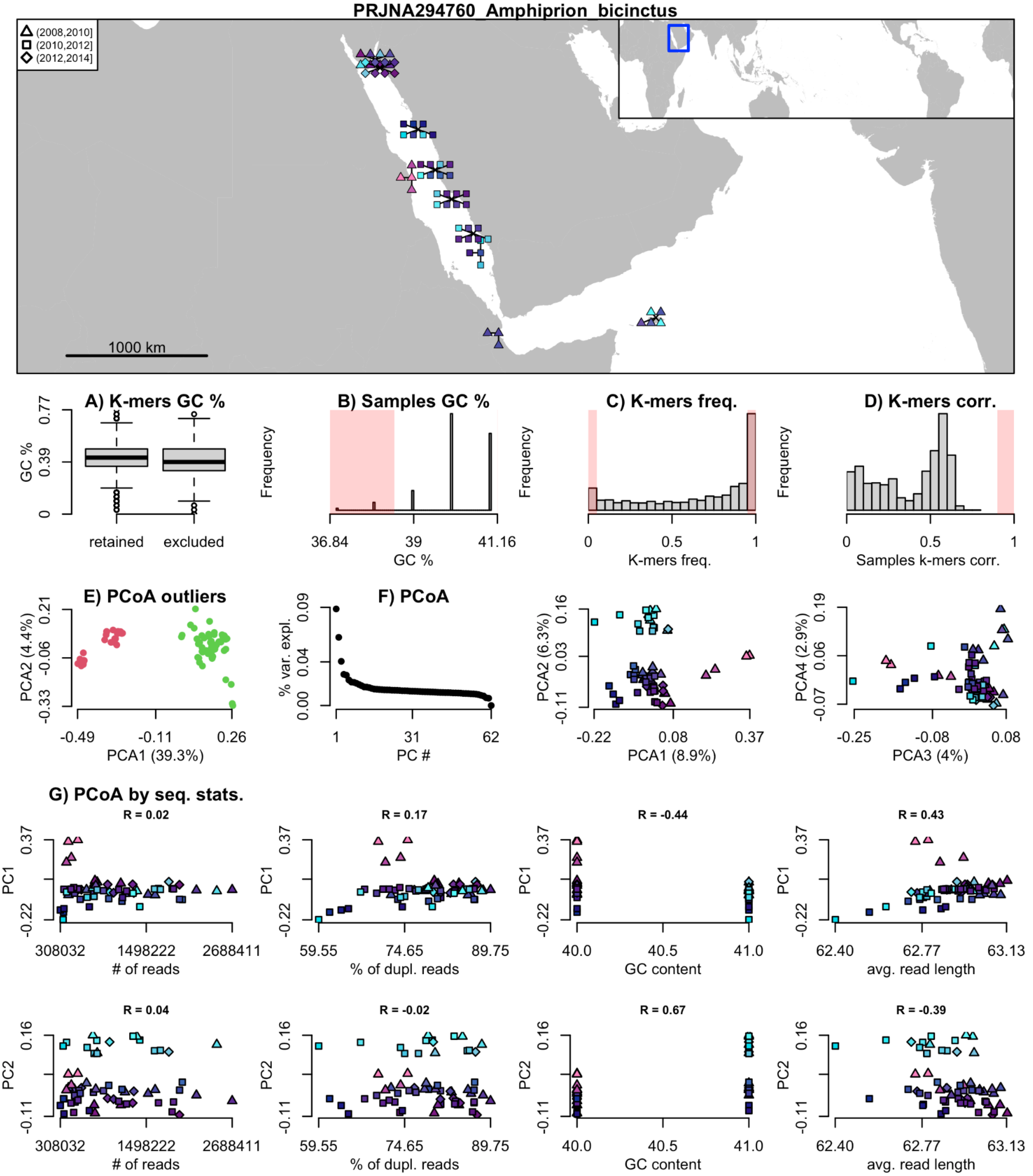

Figure 2 (continue)

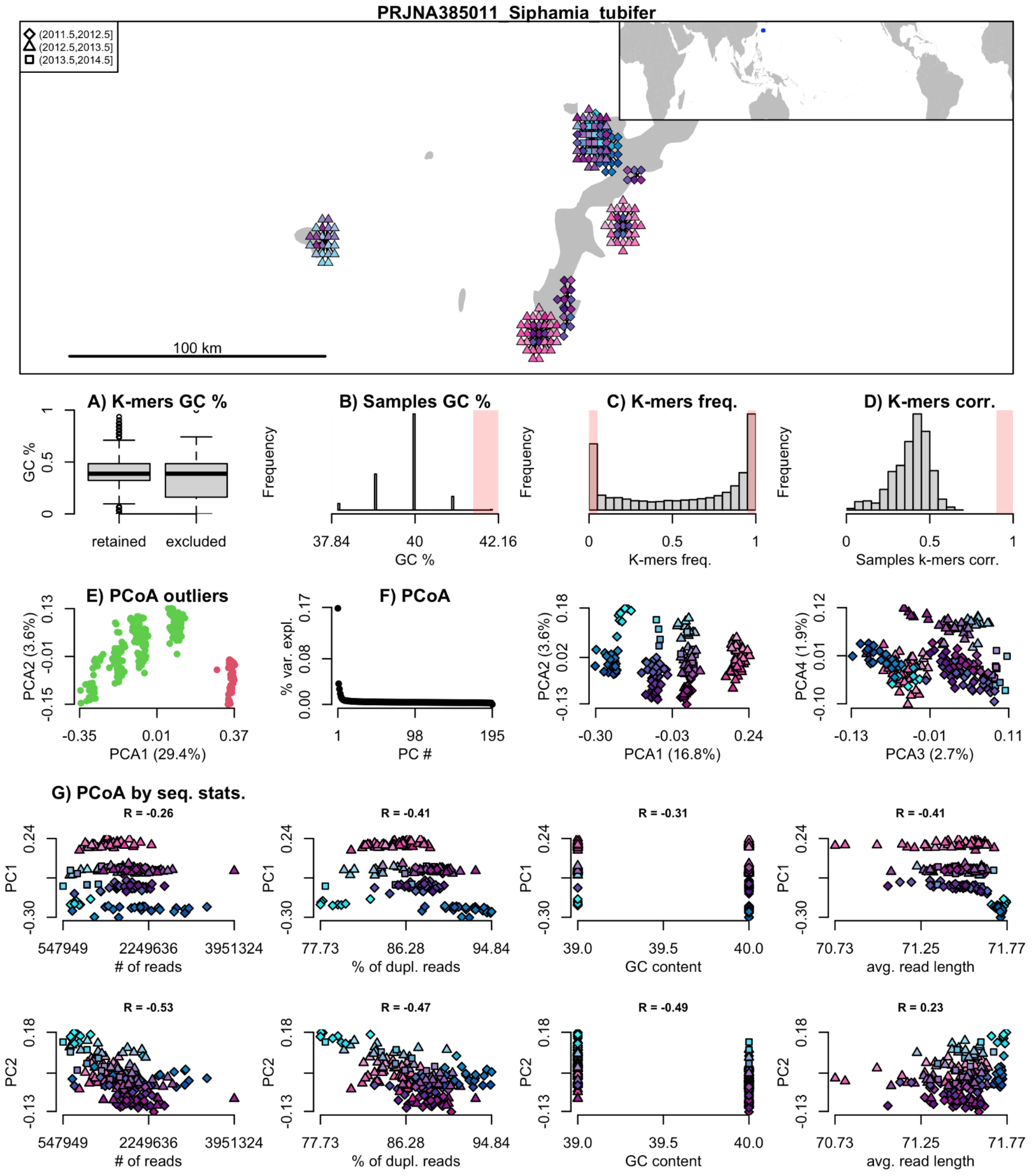

**Figure 2 (continue)**

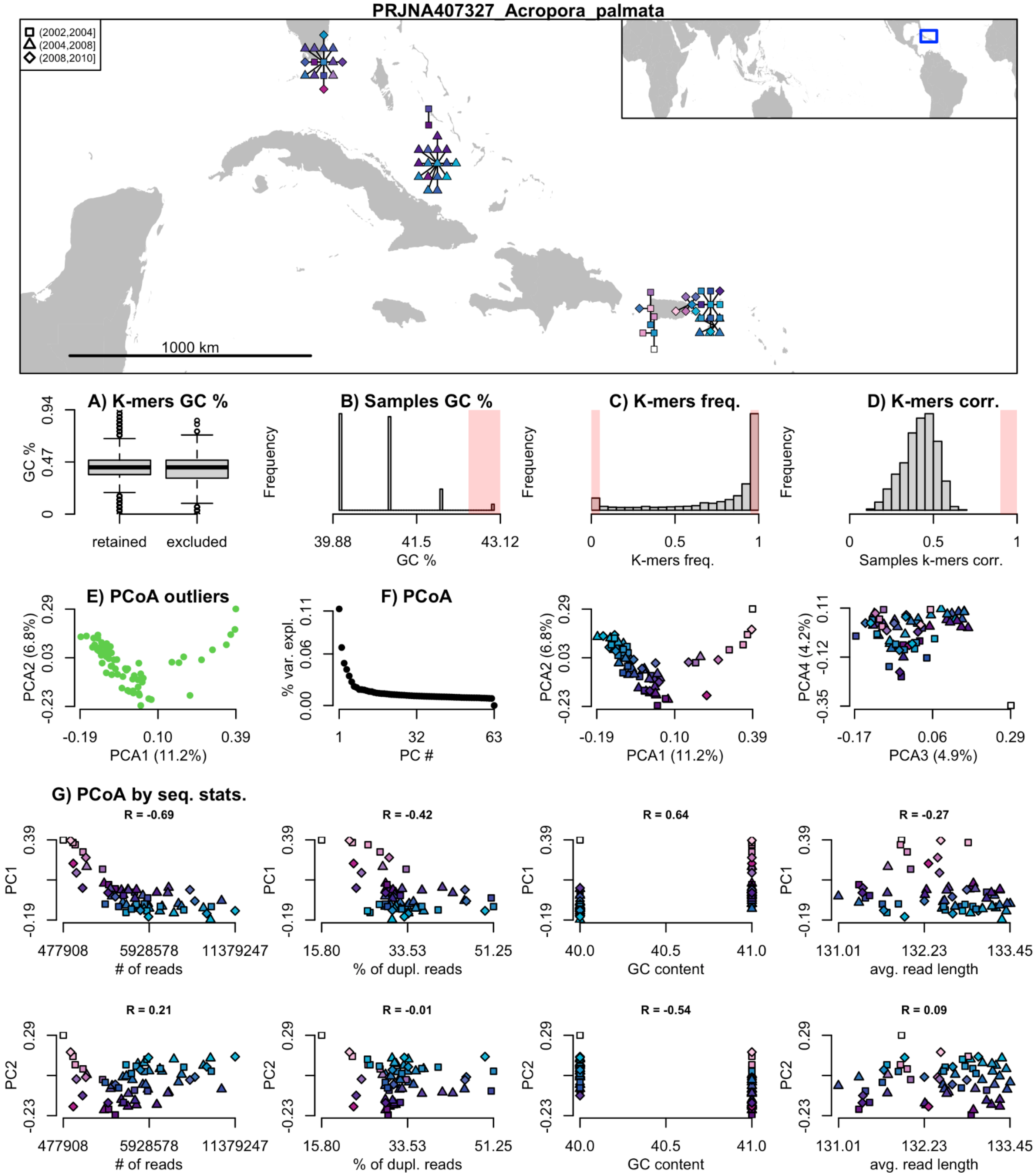

Figure 2 (continue)

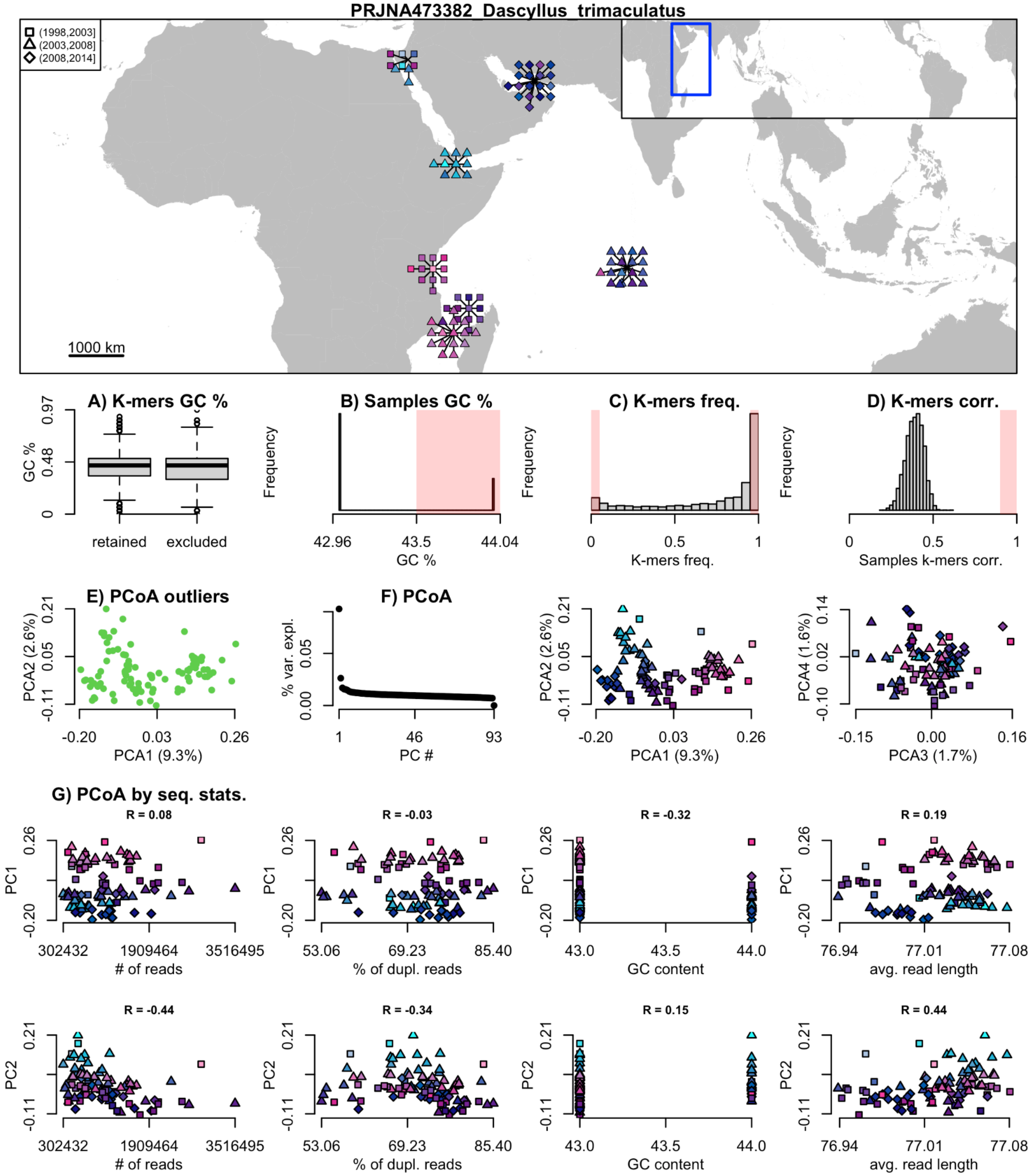

**Figure 2 (continue)**

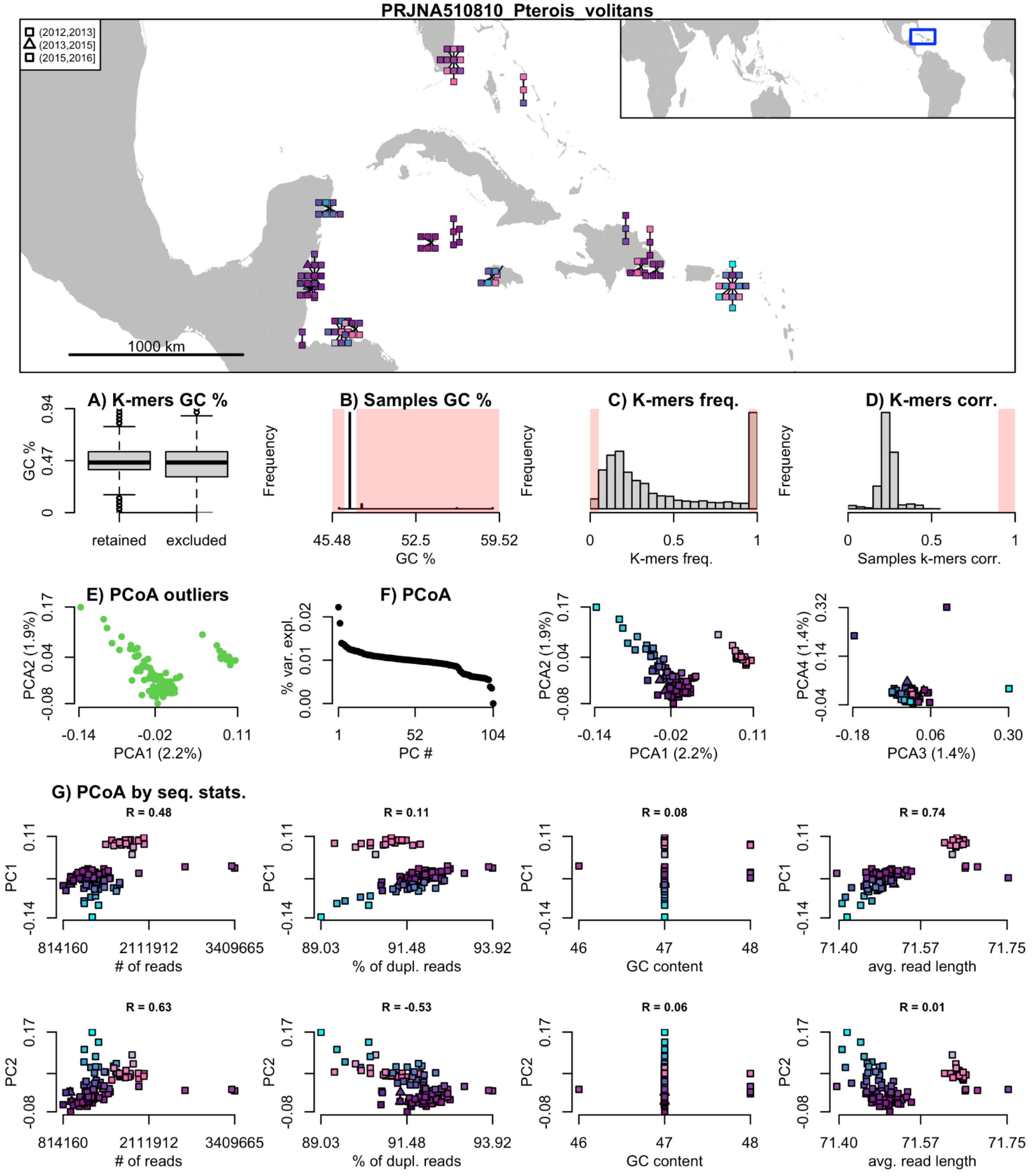

**Figure 2 (continue)**

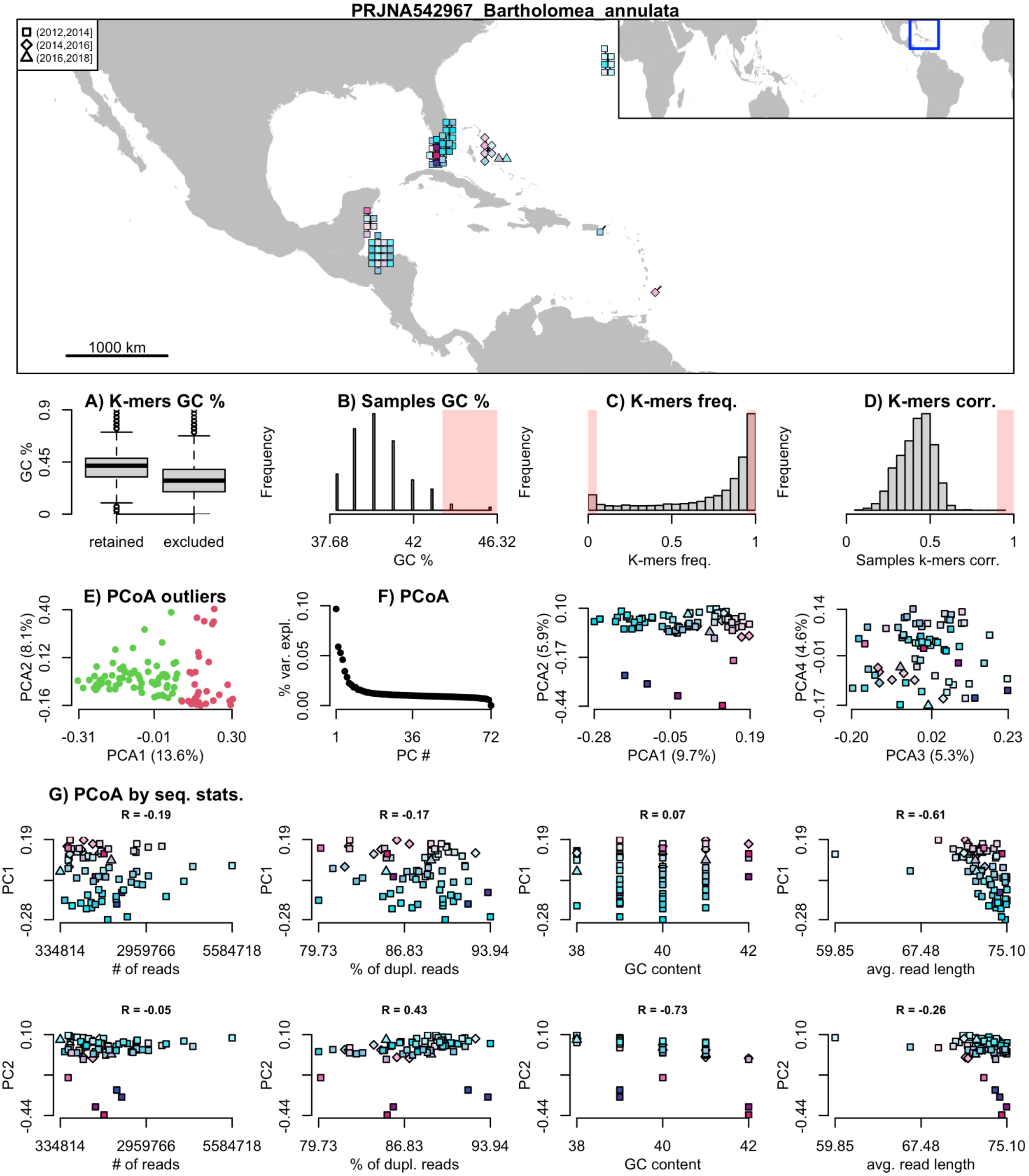

**Figure 2 (continue)**

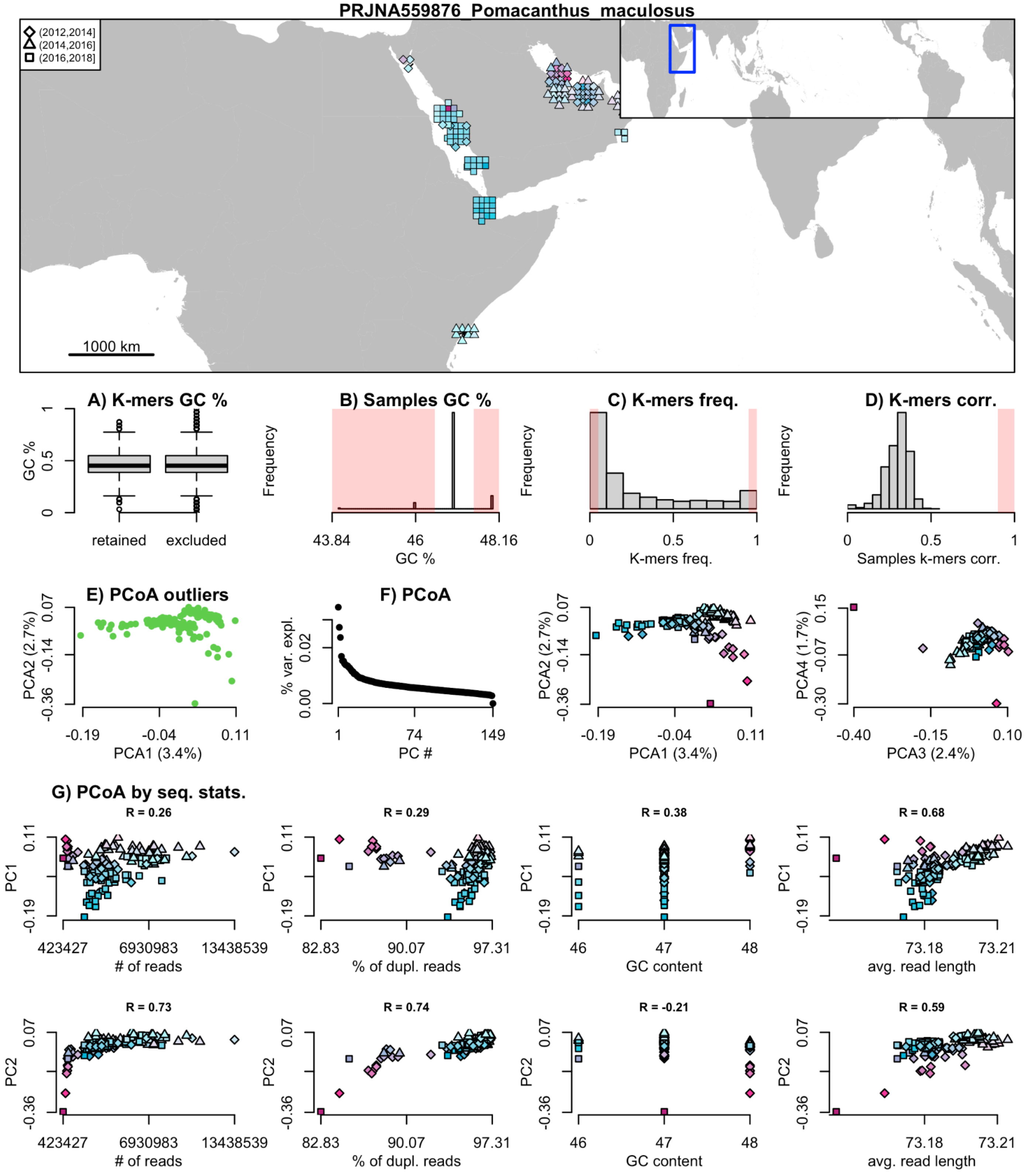

**Figure 2 (continue)**

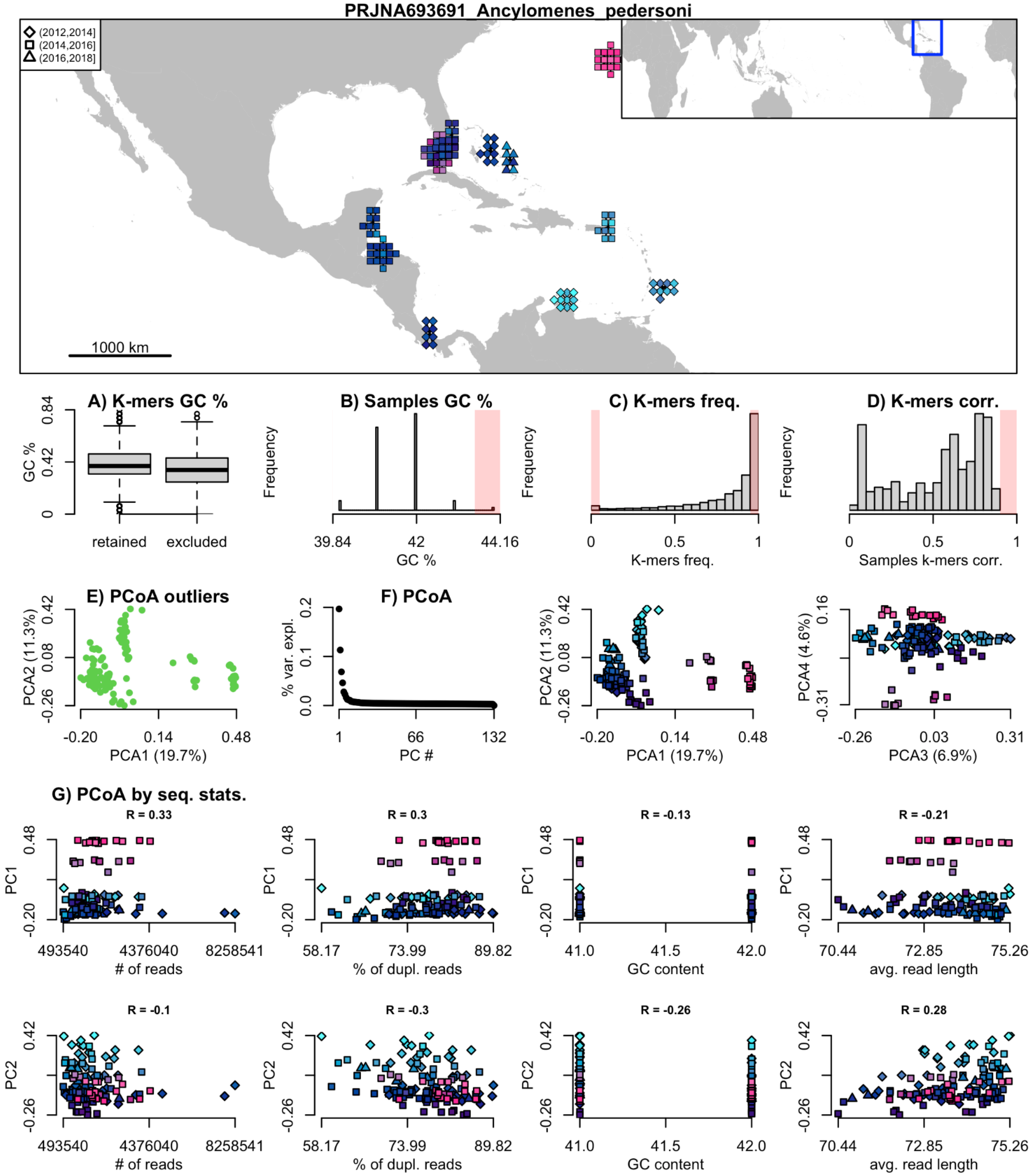

Figure 2 (continue)

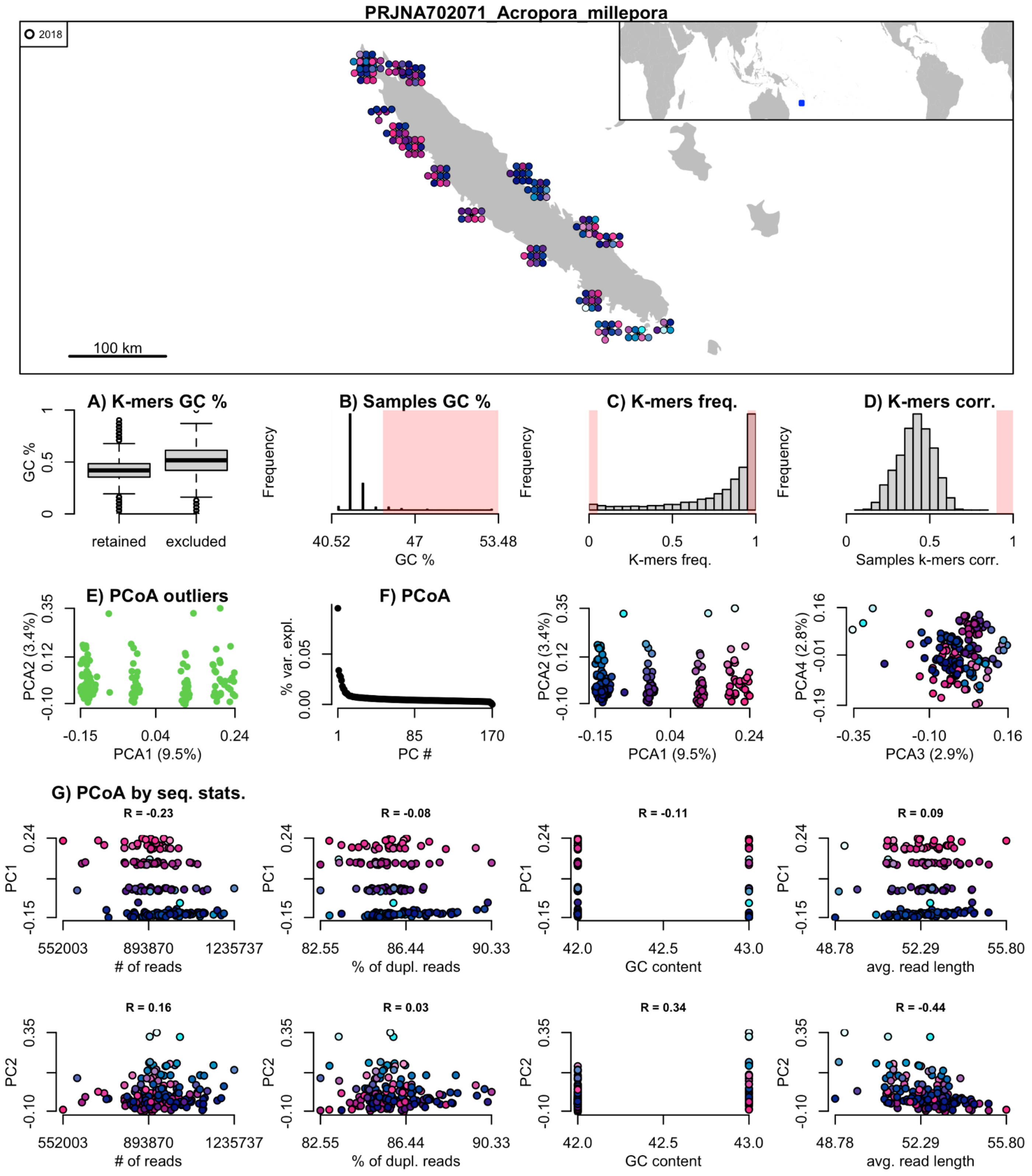

**Figure 2 (continue)**

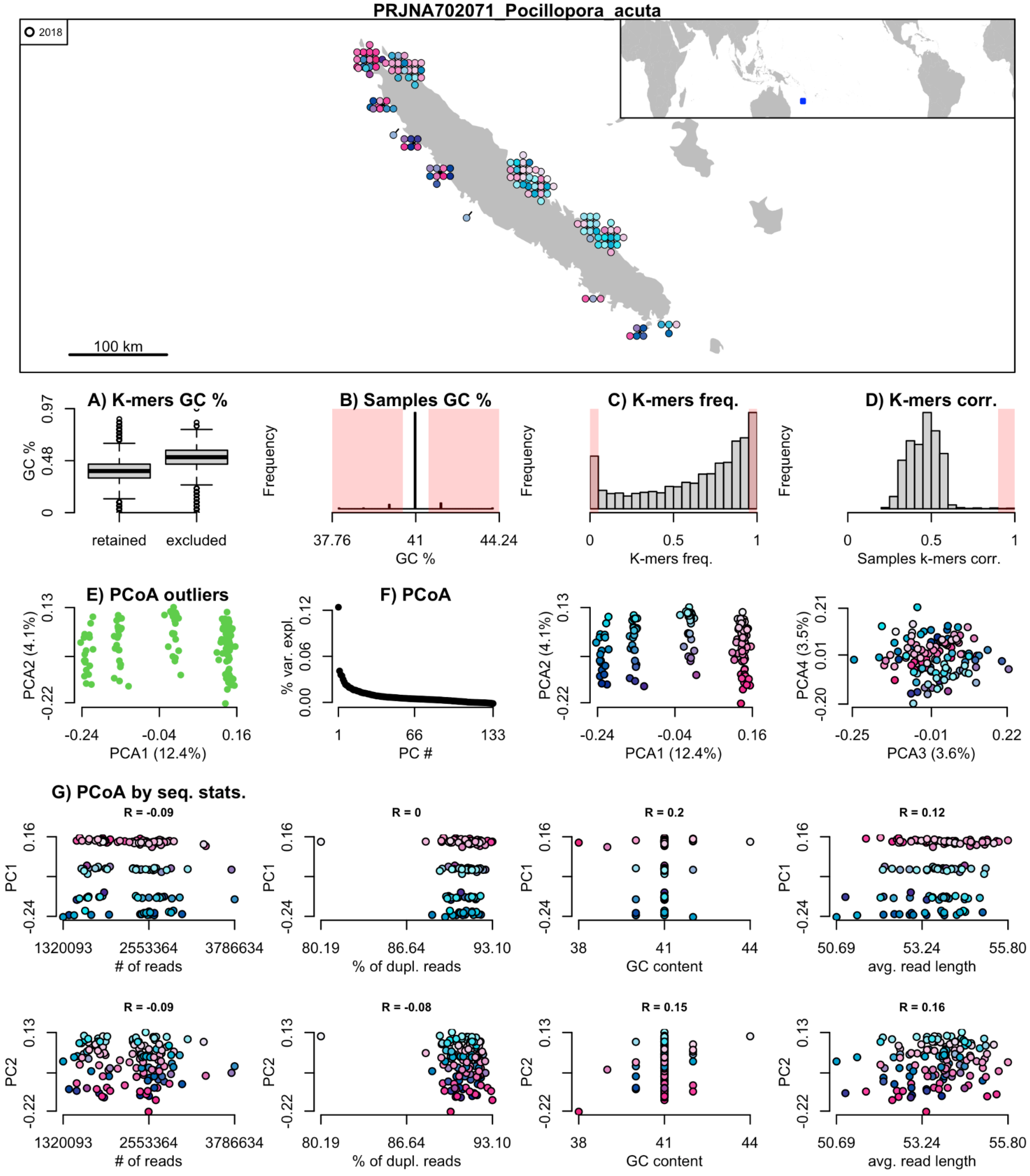

**Figure 2 (continue)**

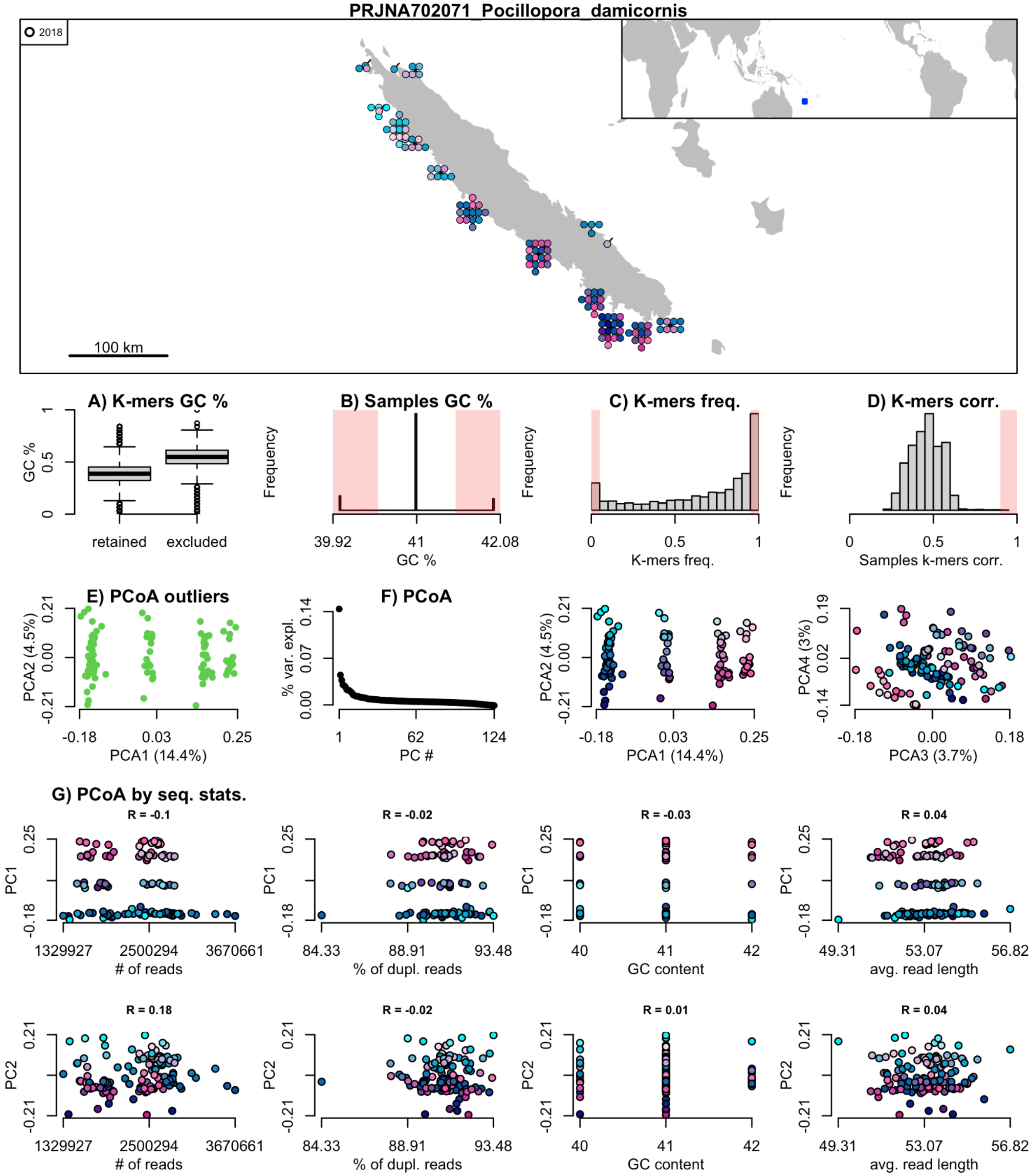

Figure 2 (continue)

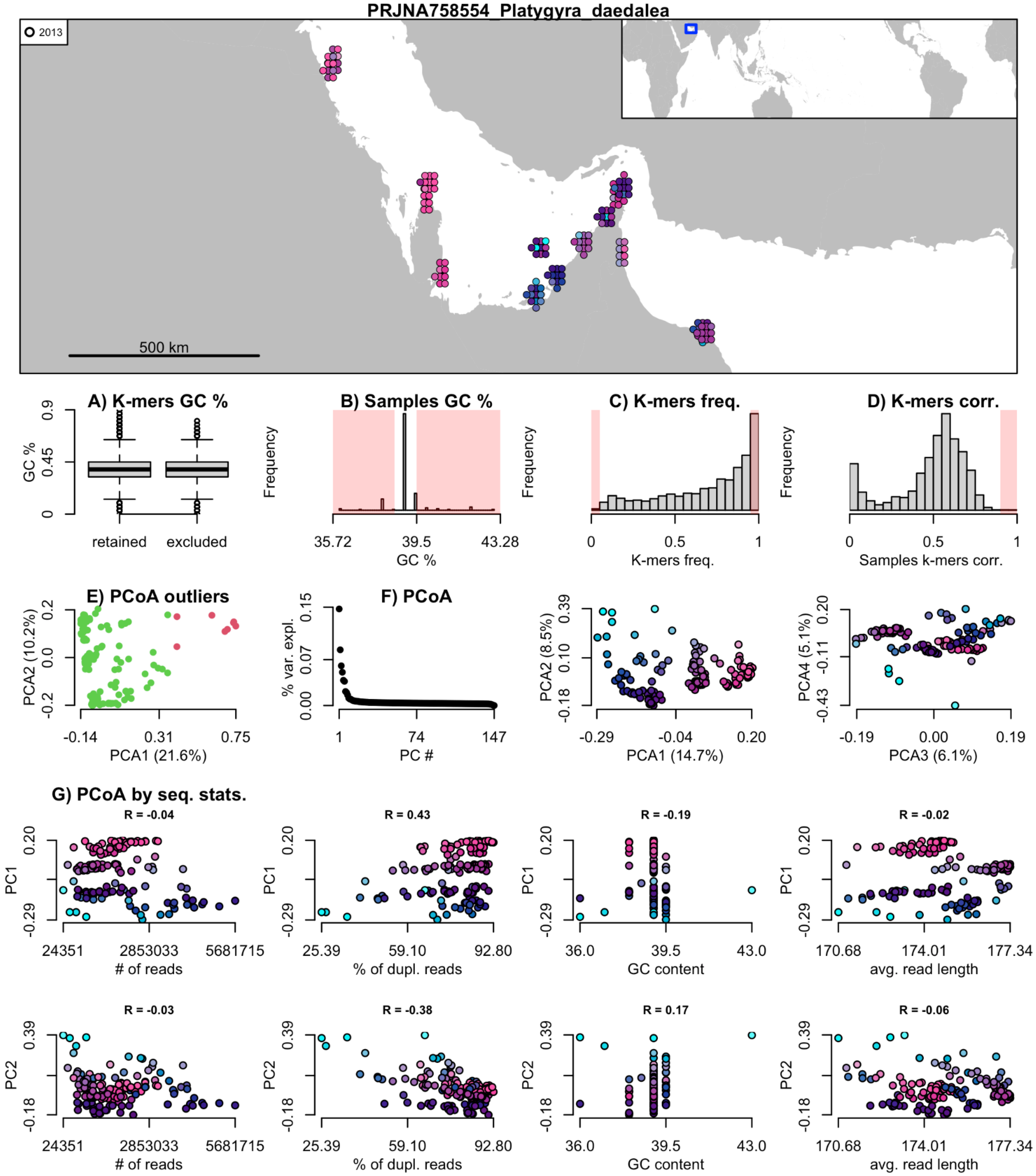

**Figure 2 (continue)**

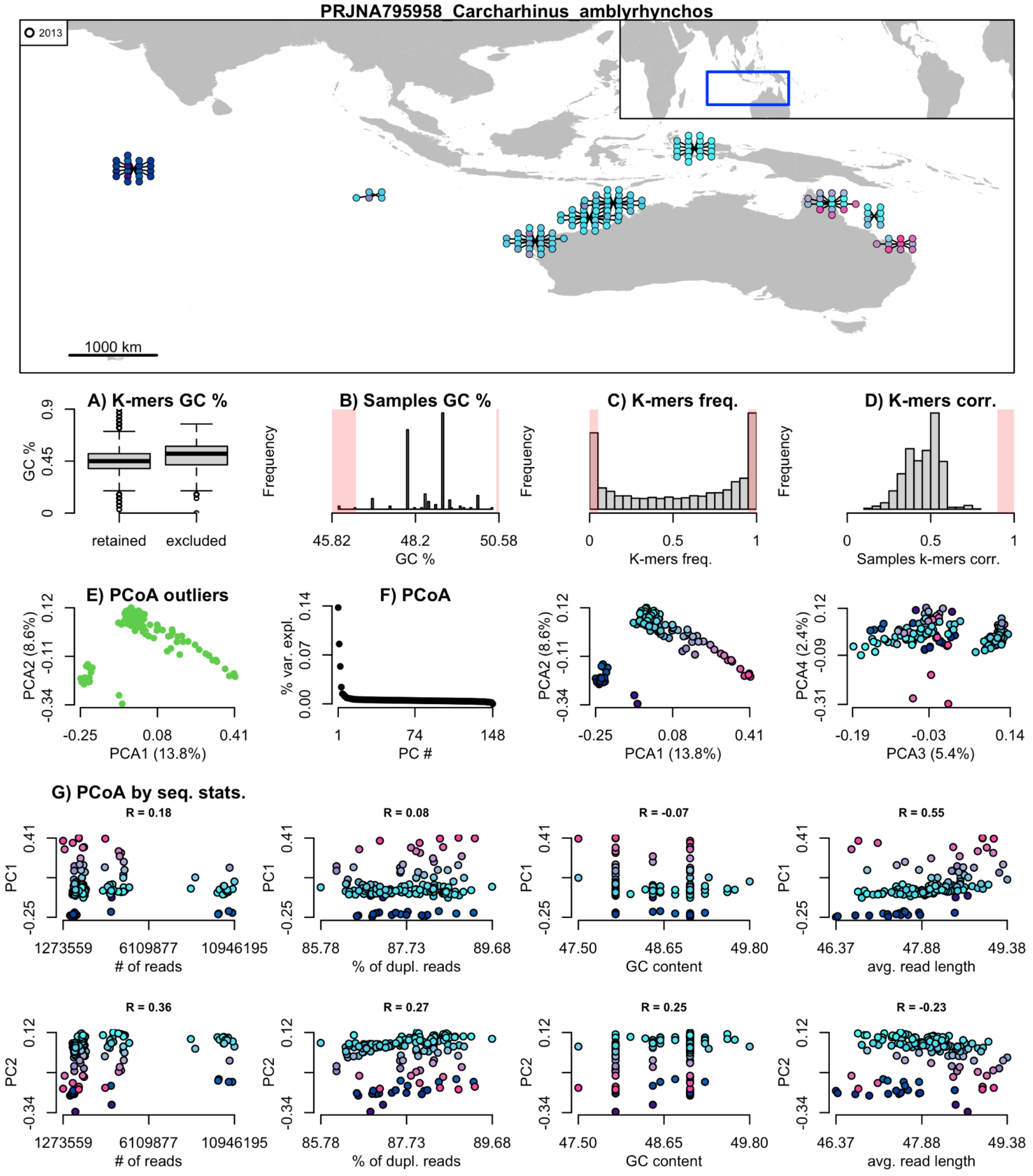

**Figure 2 (continue)**

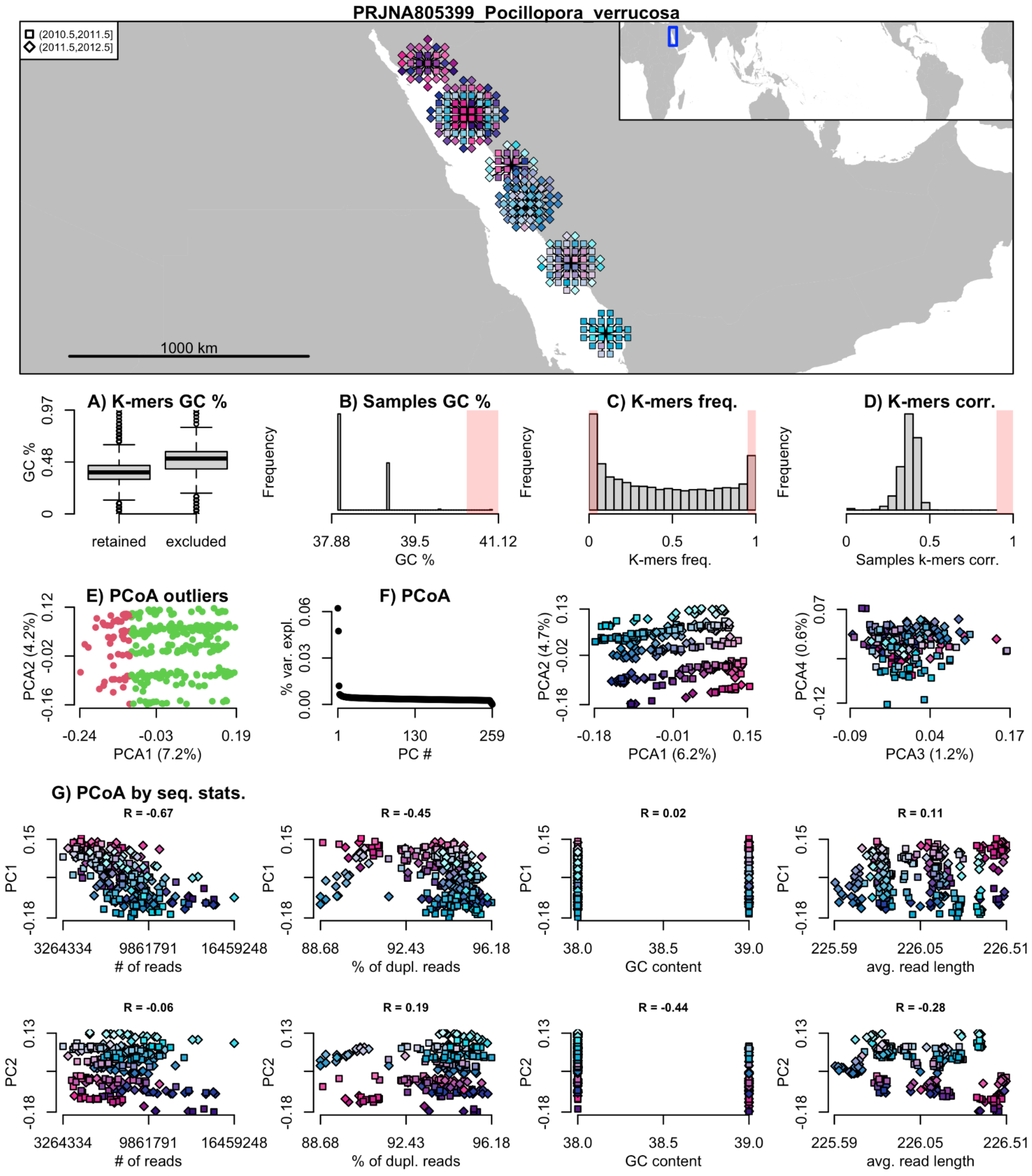

**Figure 2 (continue)**

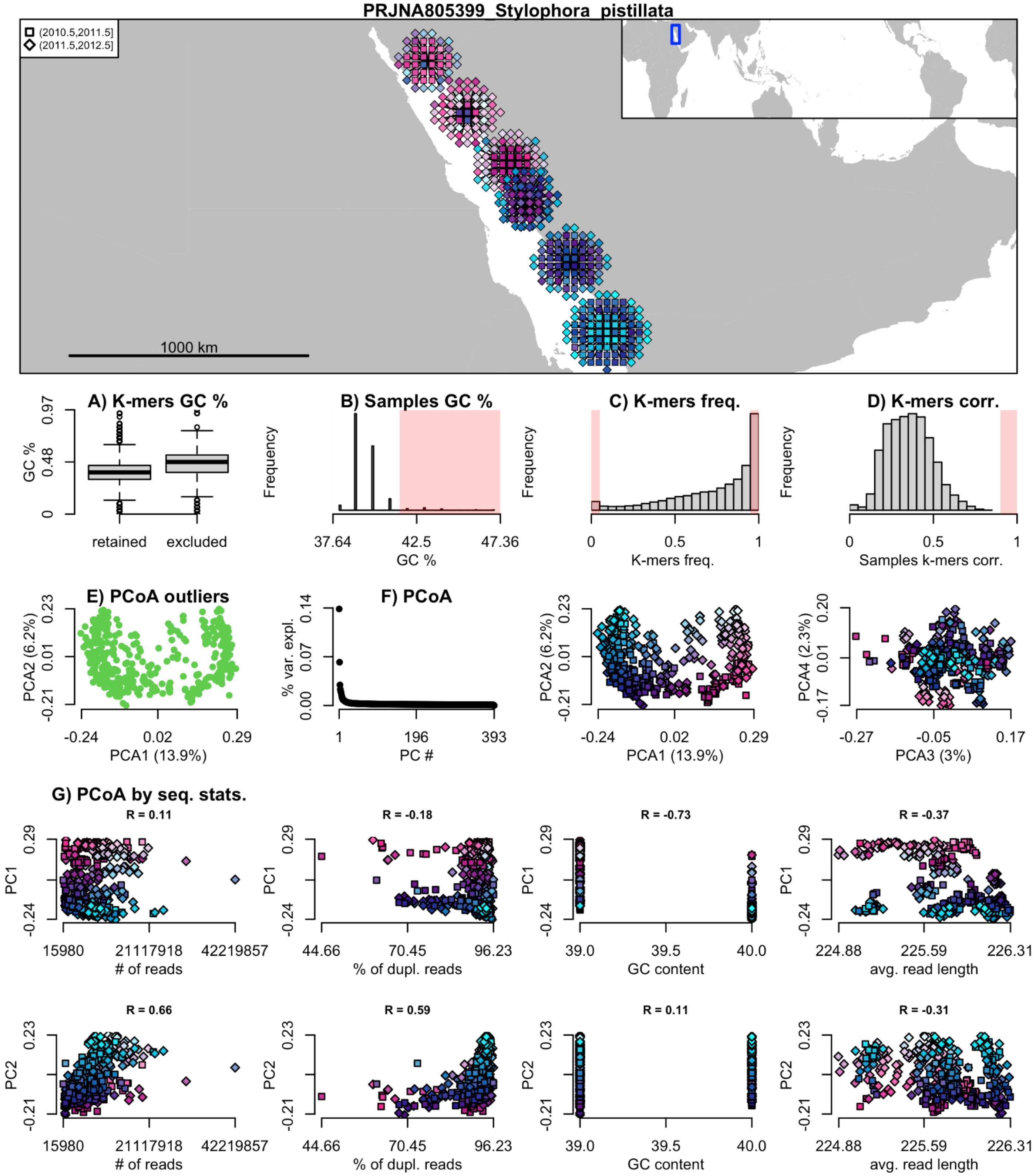

**Figure 2 (continue)**

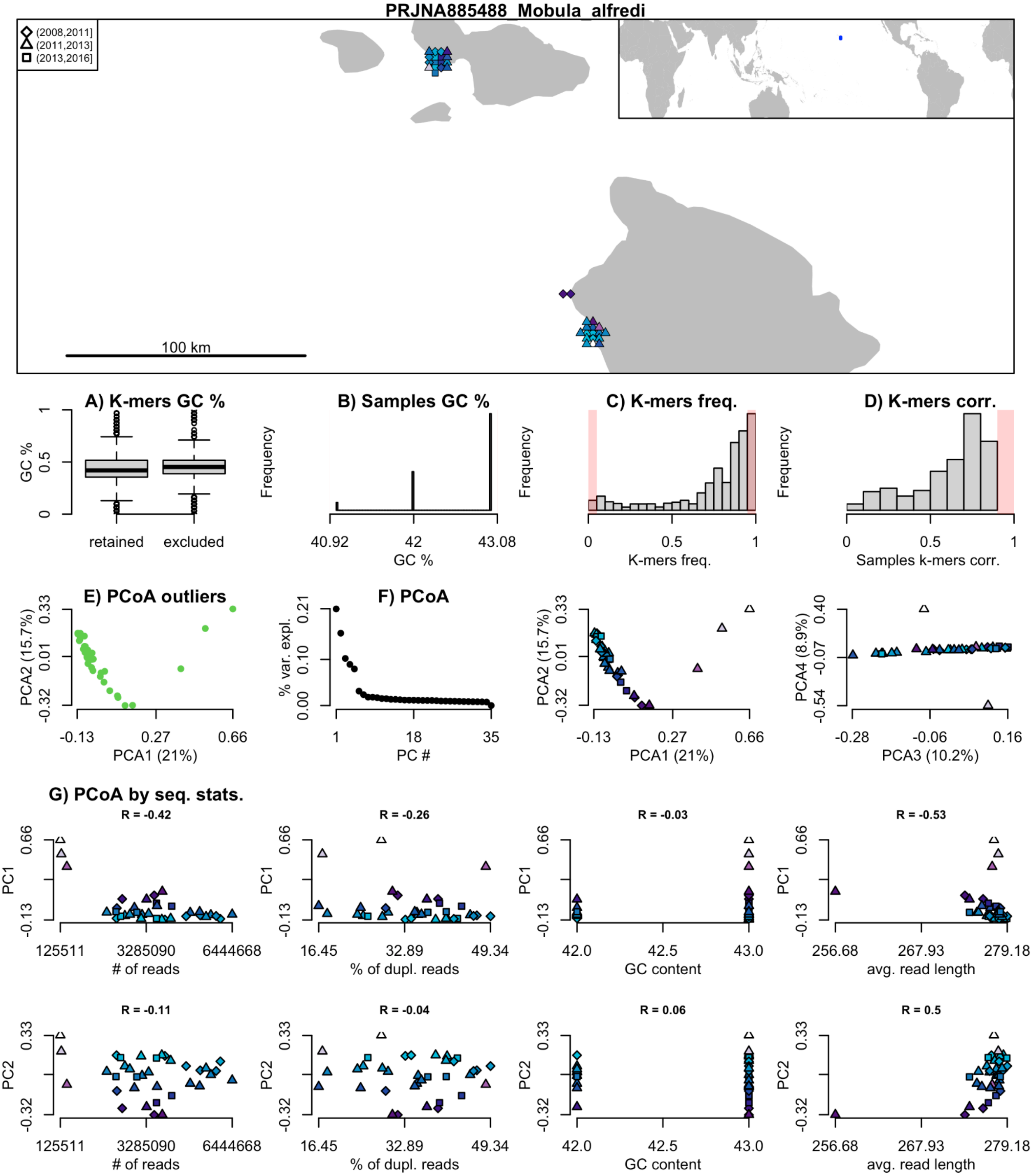

**Figure 2 (continue)**

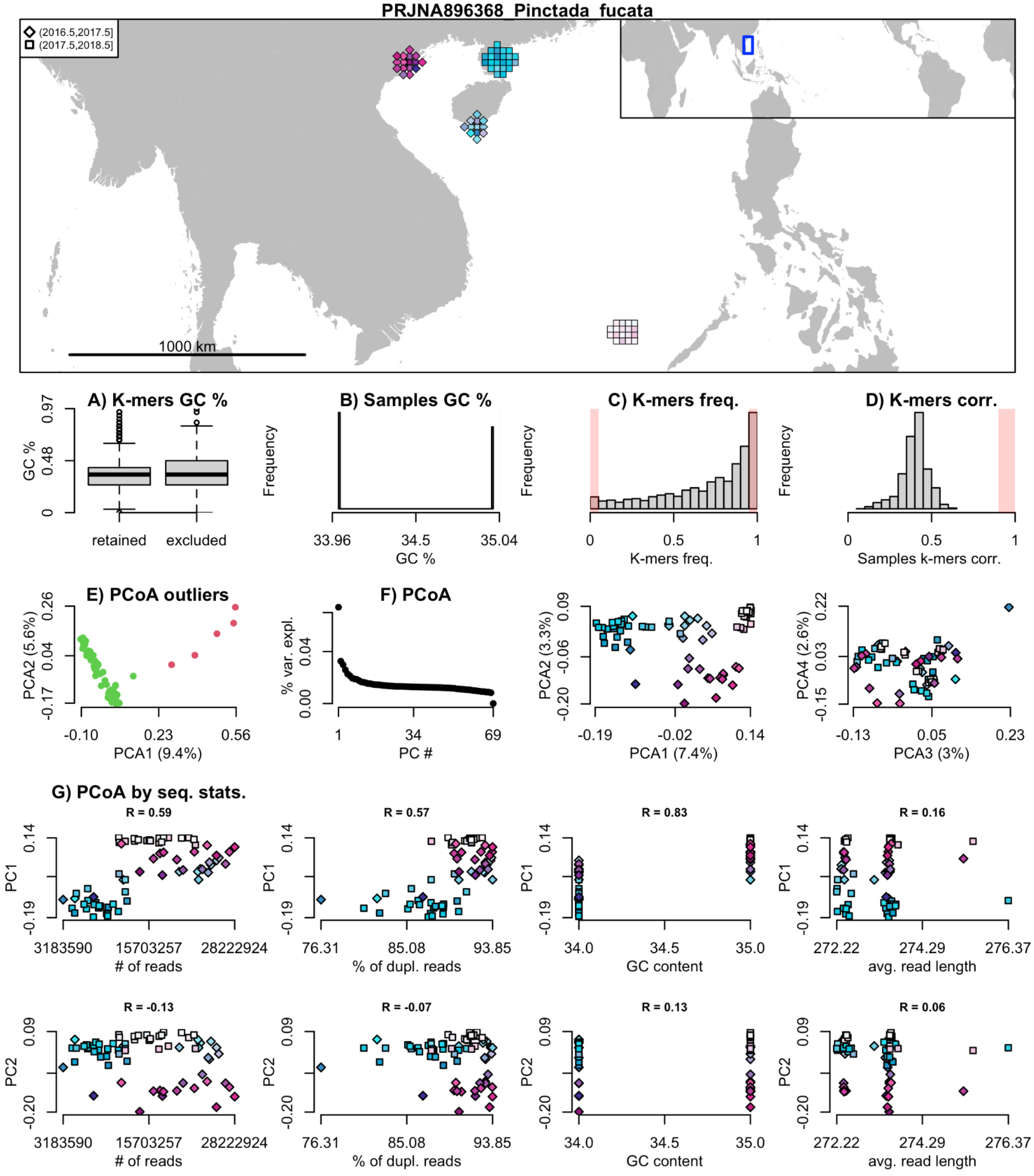

**Figure 2 (continue)**

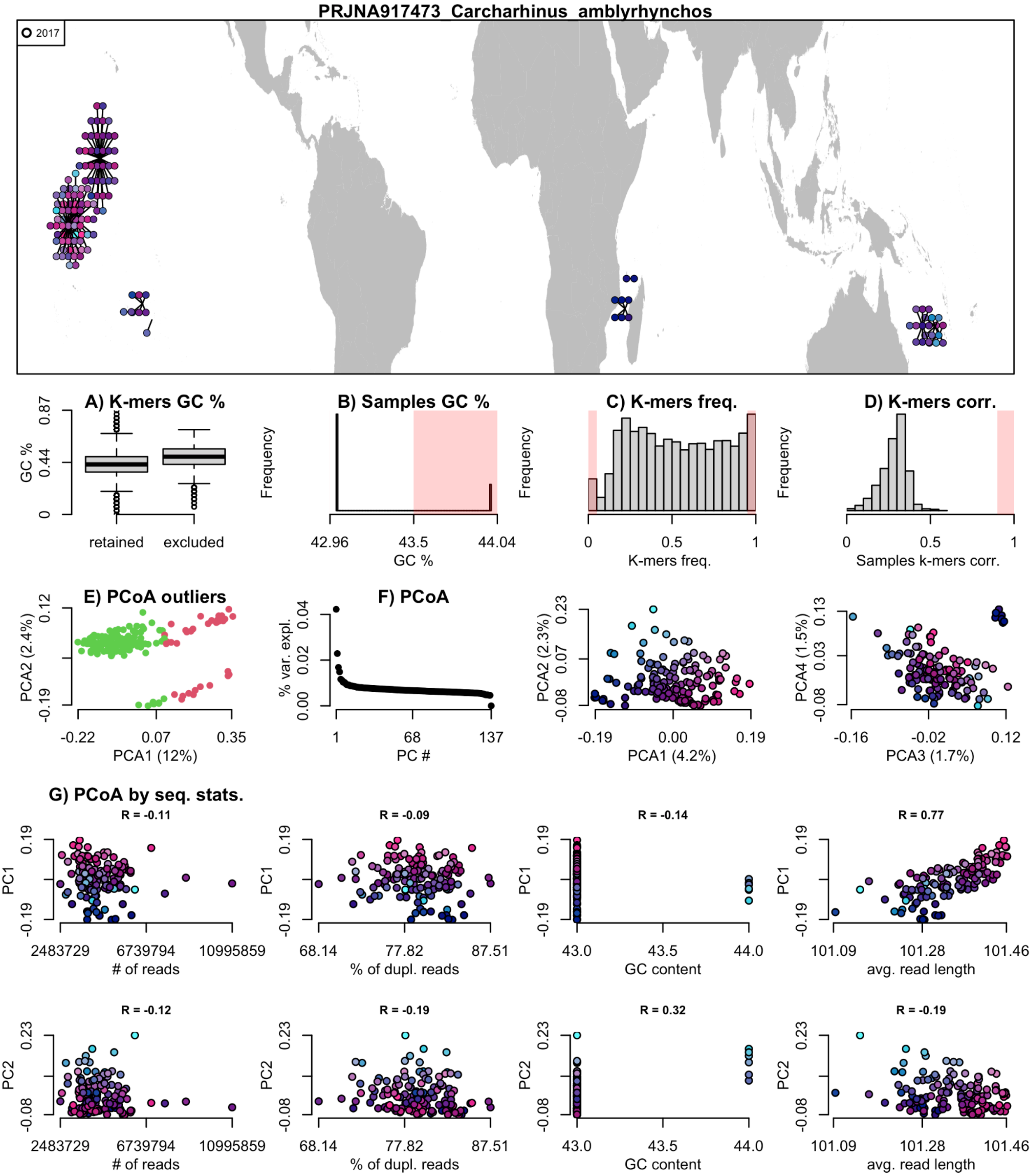

**Supplementary Figure 4. Geographic effects on genetic distances.**

The four maps show the effects of geographic grouping on Bray–Curtis genetic distances at different spatial levels: (A) ocean, (B) ecoregion, (C) sampling region (groups of sites within 100 km), and (D) sampling reef (groups of sites within 10 km). Points indicate the effects on genetic distances for sample pairs collected within the same geographic unit (e.g., same ocean), while lines indicate the effects for pairs collected across different units (e.g., between the Pacific and Indian Oceans).

**A) Effect of ocean**

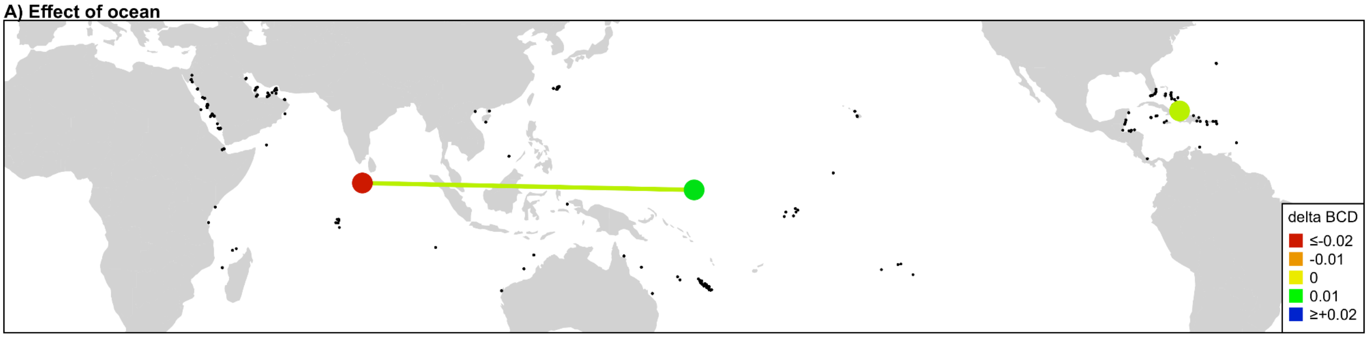


**B) Effect of ecoregion**

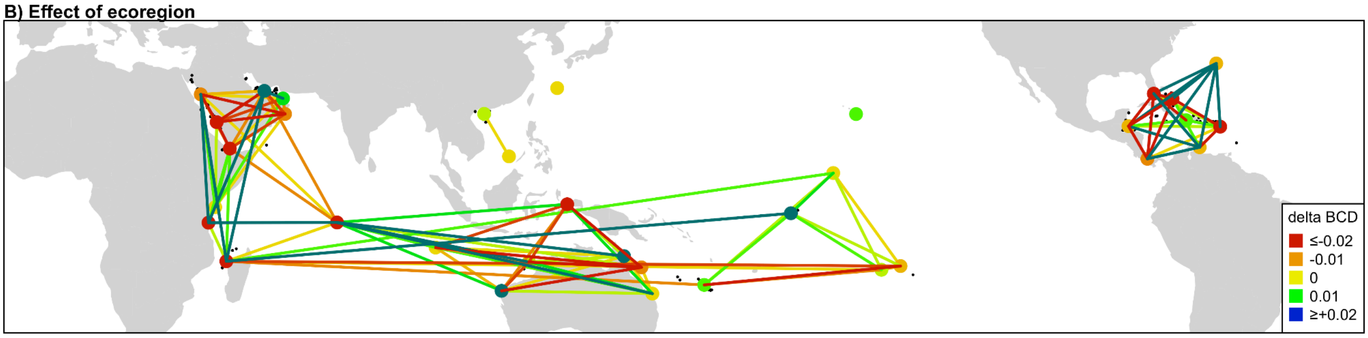


**C) Effect of sampling region**

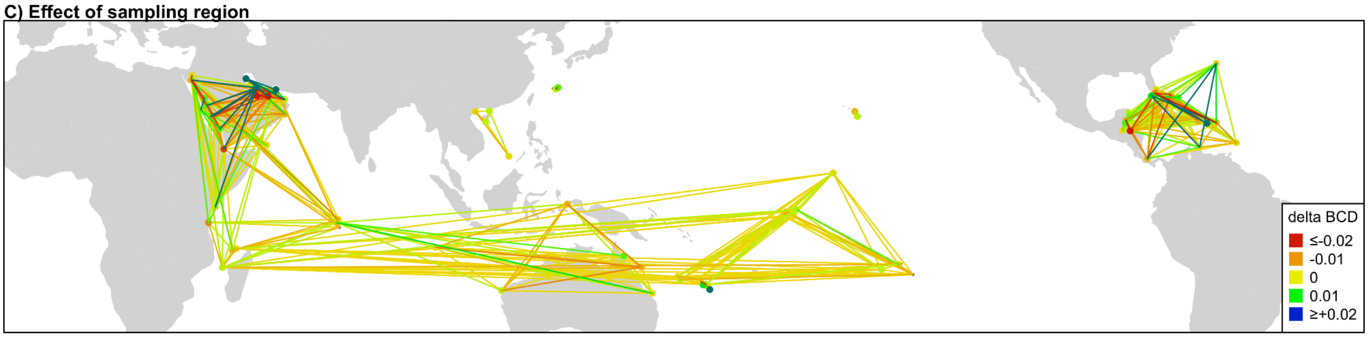


**D) Effect of sampling reef**

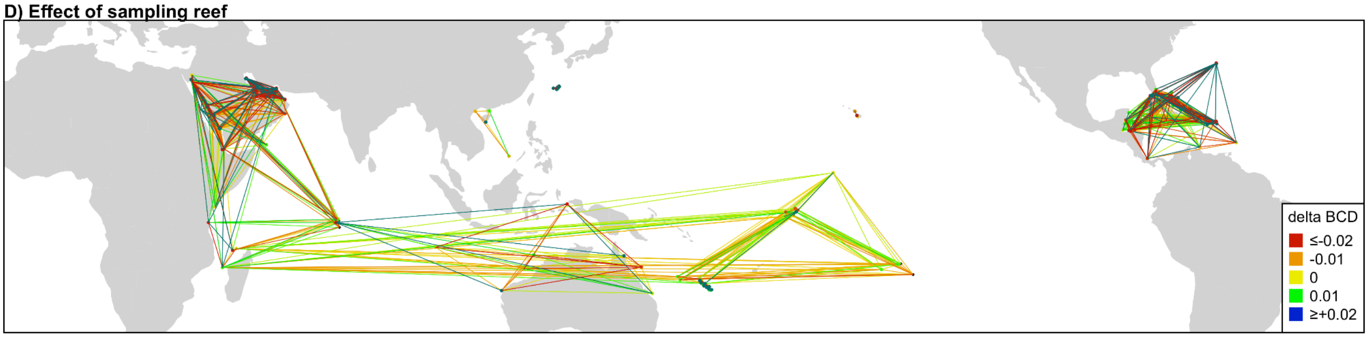

**Supplementary Figure 5. Leave-one-region-out cross-validation of logistic elastic net regularization models.**

Cross-validation results of penalized logistic regression models used to identify environmental variables predicting local effects on Bray–Curtis genetic distances. Each panel corresponds to a leave-one-region-out cross-validation at different spatial scales: (A) ecoregion (~800 km), (B) Large Marine Ecosystem (~1,500 km), and (C) oceanic realm (~5,000 km). For each scale, shown are: a map indicating reef groupings by region, a boxplot of AUC values across alpha levels (optimal alpha highlighted in purple), AUC variation across lambda values at optimal alpha (optimal lambda in purple), and the AUC curve under optimal alpha and lambda.

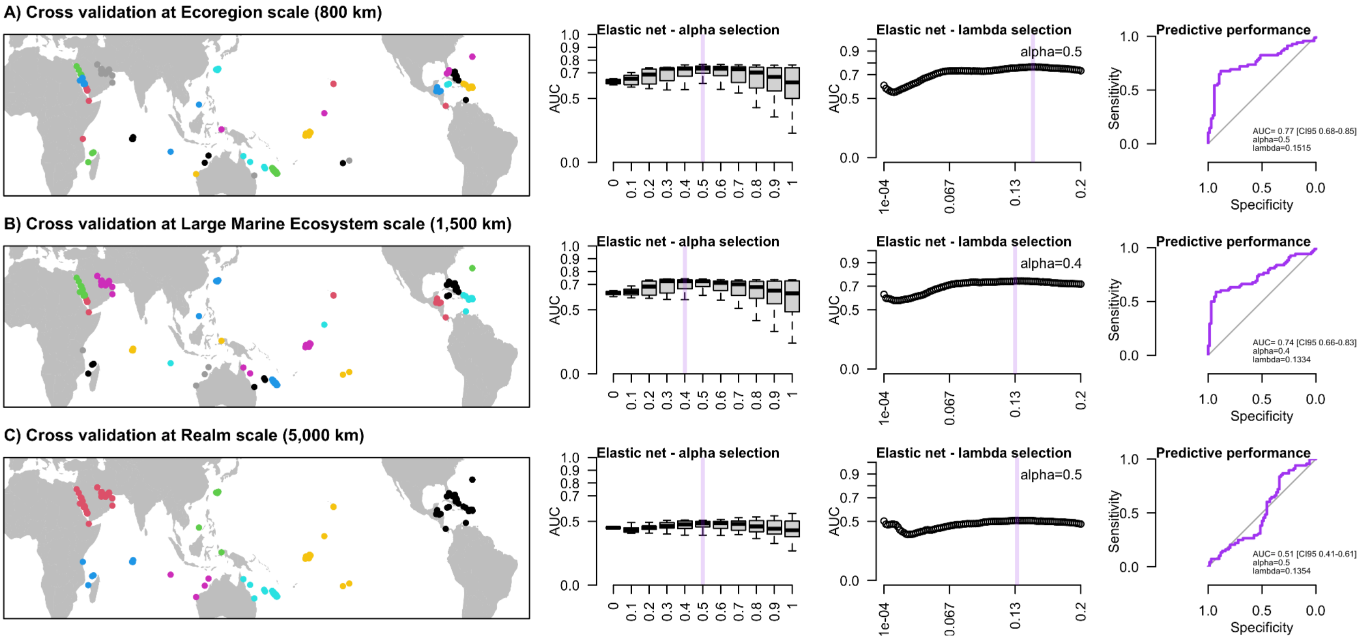

**Supplementary Figure 6. Leave-one-region-out cross-validation of linear elastic net regularization models.**

Cross-validation results of penalized linear regression models used to identify environmental variables predicting local effects on Bray–Curtis genetic distances (BCD). The analysis employed a leave-one-region-out approach, with mean absolute error (MAE) between real and predicted values as the performance metric. (A) shows MAE variation when performing the cross-validation on regions covering different spatial scales: ecoregion (grouping reefs within ~800 km), Large Marine Ecosystem (LME, ~1,500 km), and oceanic realm (~5,000 km). The purple vertical line marks the scale with the lowest MAE (LME). (B) and (C) display MAE variation with different tuning parameters for the penalized regression: alpha (B) and lambda (C). The optimal values minimizing MAE are indicated by purple vertical lines. (D) compares predicted vs. real BCD for the best model (optimal spatial scale, alpha, and lambda). (E) shows the MAE for the optimal model from (D; “Environmental prediction”) in comparison to the MAE obtained with an analogous model built on spatial distances (distance based Moran Eigenvector Maps) instead of environmental variables.

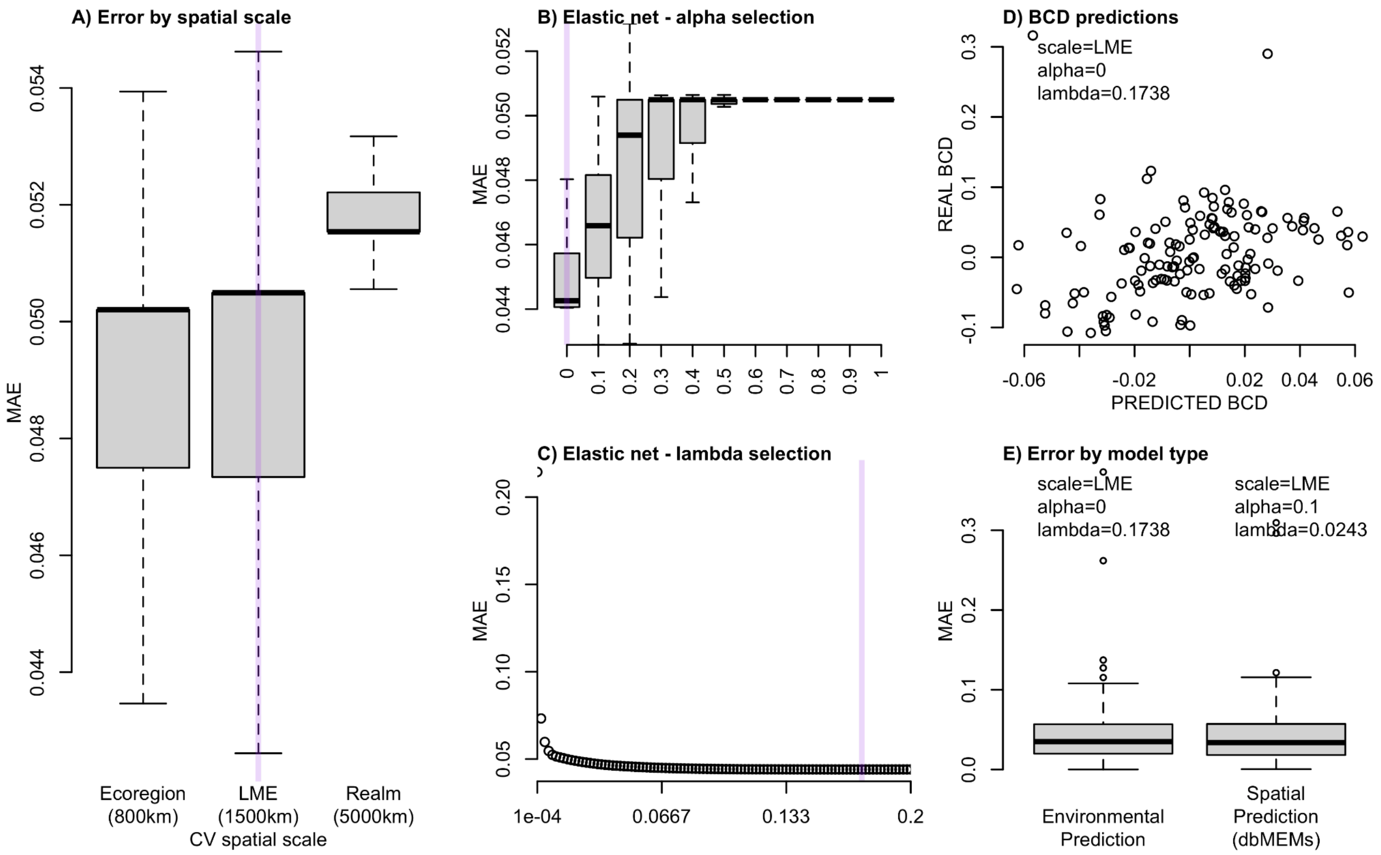

**Supplementary Figure 7. Elastic net regularization model using spatial predictors.**

Cross-validation results of a penalized logistic regression model using spatial predictors (Moran Eigenvector Maps, MEMs) to predict local effects on Bray–Curtis genetic distances. (A) Variation in Area Under the Curve (AUC) across alpha values; optimal alpha is highlighted in purple. (B) AUC variation across lambda values under the optimal alpha; optimal lambda is highlighted. (C) AUC under optimal alpha and lambda settings. (D) Spatial distribution of MEMs retained in the final model.

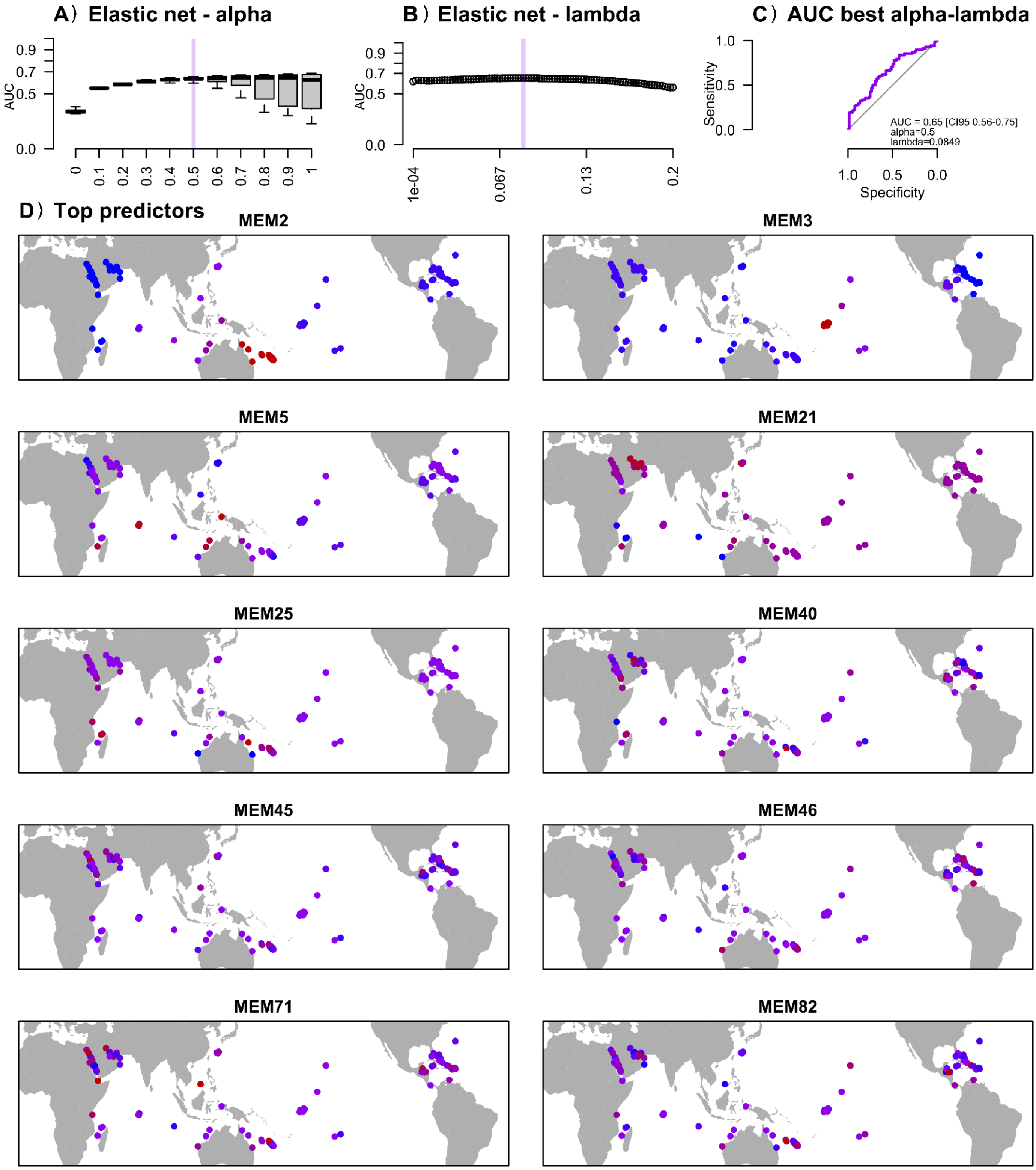

**Supplementary Figure 8. Spatial and temporal distribution of datasets.**

For each of the 19 datasets included in the analysis (color-coded), the top panel shows the spatial distribution of sampling locations, with lines connecting reefs sampled within the same dataset. The bottom panel displays the spatial (x-axis: longitude) and temporal (y-axis: year of sampling) distribution of sampling locations, with lines connecting sites sampled in the same dataset.

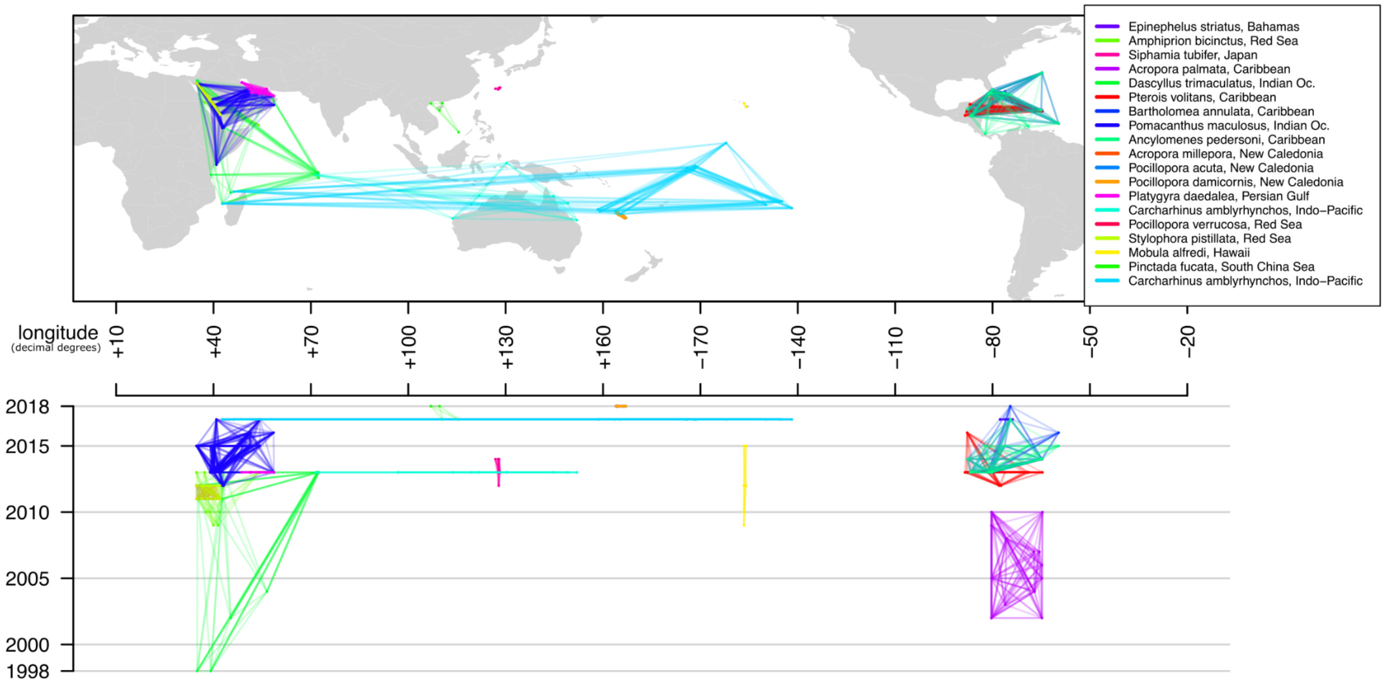

**Supplementary Figure 9. Unadjusted associations between genetic distances and sampling design variables.**

The figure shows associations of Bray-Curtis genetic distances (BCD) between sample pairs with the following variables: (A) Dataset, (B) geographic distance between sampling sites, (C) temporal distance between sampling years, (D) mid-point of the sampling years, (E) absolute mid-point latitude of the sampling sites. For panels (C–E), associations are presented for all sampling pairs ("all geo distance"), as well as for pairs separated by less than 1 km, 1–10 km, 10–100 km, or more than 100 km. In panels (B–E), lines indicate the direction of Pearson correlation coefficients. Note: These associations are unadjusted and reflect BCD relationships with each variable independently, without accounting for other variables.

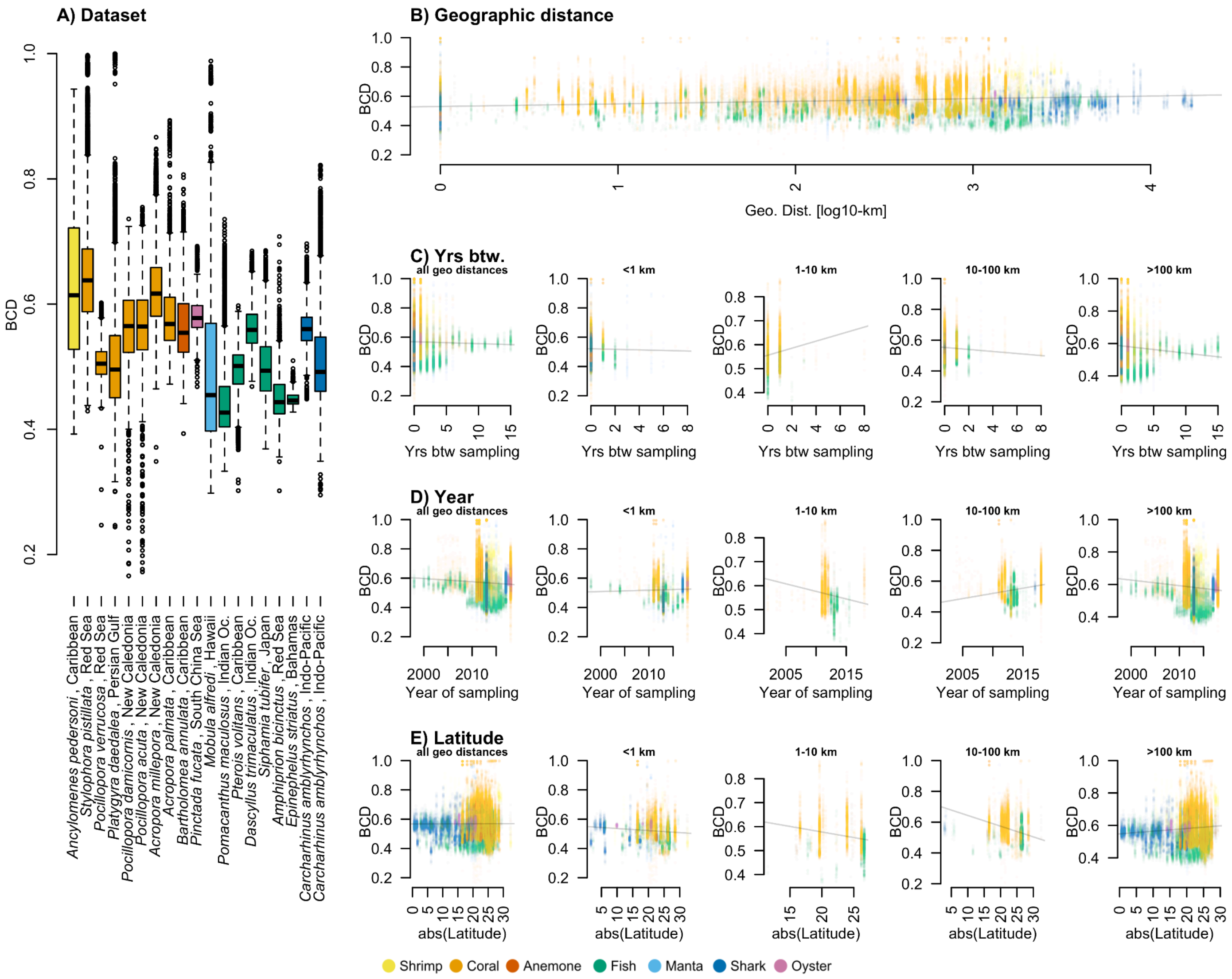

**Supplementary Figure 10. Dataset-specific effects on genetic distances.**

The three panels show how predicted Bray–Curtis genetic distances vary across datasets for different predictors: (A) aerial distance between sample pairs, (B) number of years between sampling, (C) midpoint year of sampling. Colors represent different taxonomic groups (see legend below).

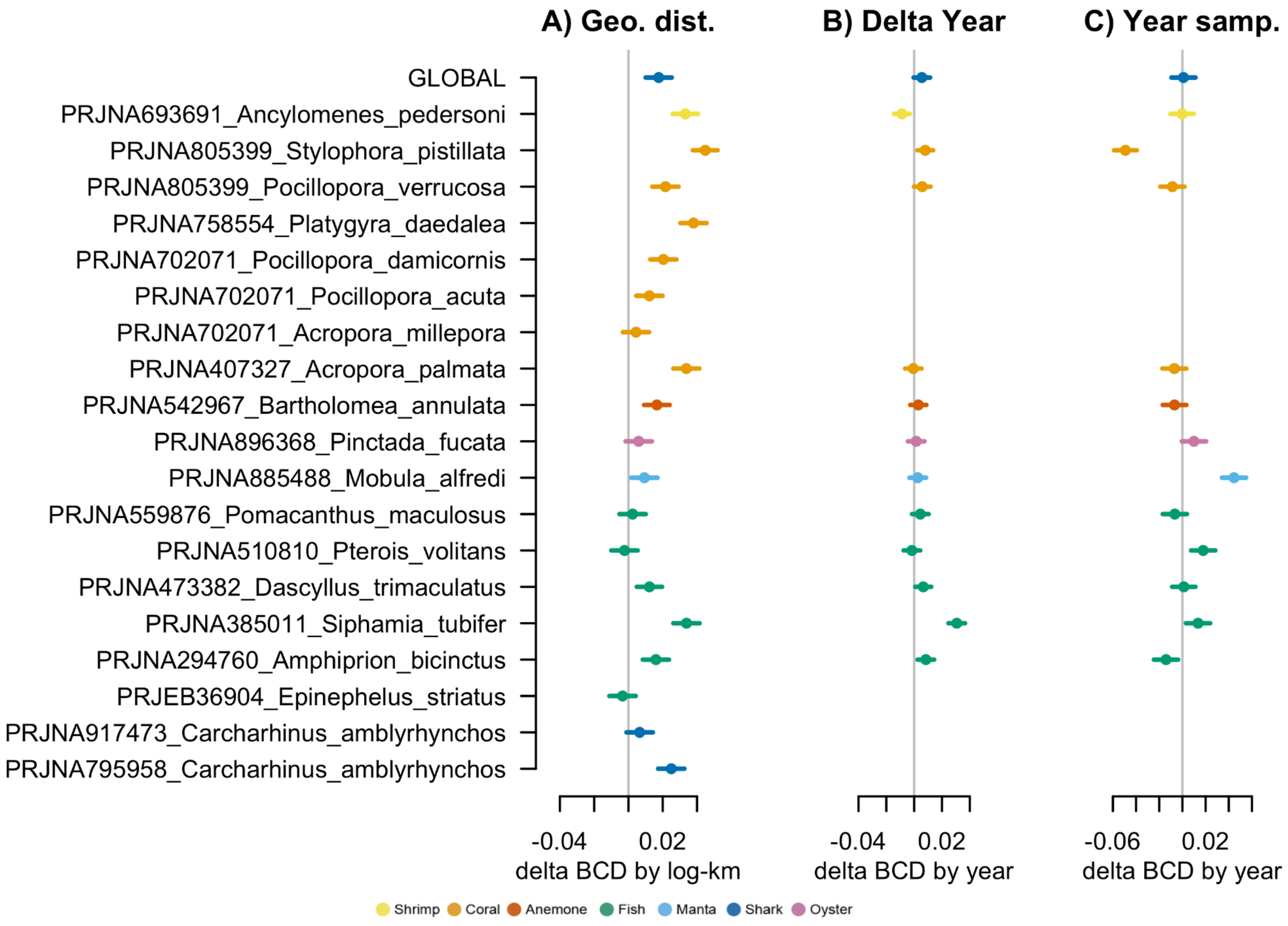

**Supplementary Figure 11. Effect of the interaction between aerial and temporal distance on genetic distances.**

Each panel shows sample pairs collected over a different aerial distance range (<1 km, 1–10 km, 10–100 km, or >100 km), and displays the association between model-adjusted Bray–Curtis genetic distances (y-axis) and the number of years between their sampling (x-axis). Regression lines with 95% confidence intervals are shown. Points represent partial residuals and are colored by taxonomic group (see legend below).

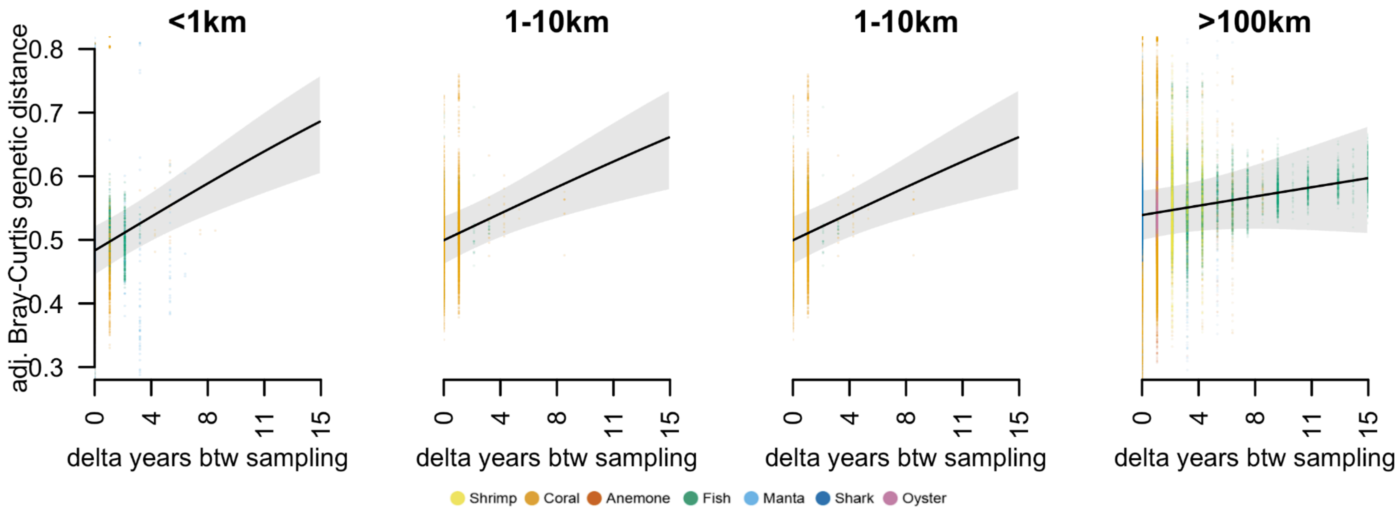

**Supplementary Figure 12. Effect of the interaction between aerial distance and latitude on genetic distances.**

Each panel represents sample pairs grouped by aerial distance: <1 km, 1–10 km, 10–100 km, and >100 km. For each group, shown is the relationship between model-adjusted Bray–Curtis genetic distances (y-axis) and the midpoint of absolute latitude (x-axis). Points represent partial residuals from the generalized linear mixed model (GLMM) and are colored by taxonomic group (see legend below). Solid lines indicate fitted GLMM regression trends, with shaded areas representing 95% confidence intervals.

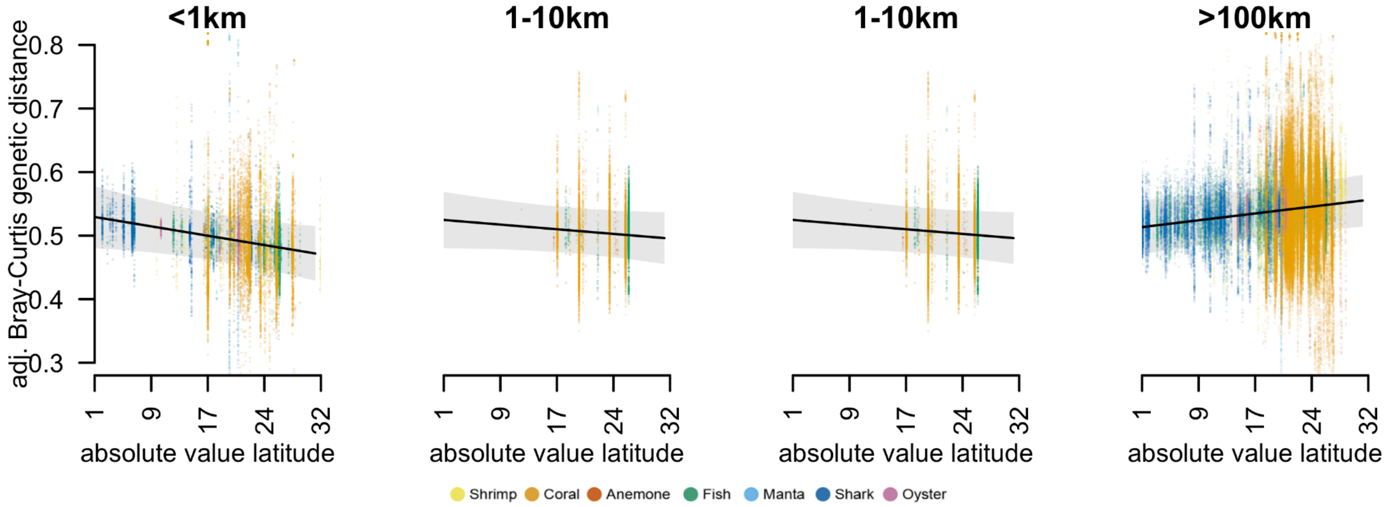

**Supplementary Figure 13. Environmental predictors retained after elastic net regularization.**

Shown are the standardized regression coefficients for environmental predictors retained in the penalized logistic regression model explaining local effects on coral reef Bray–Curtis genetic distances. All predictors were selected through elastic net regularization. For each variable, the spatial scale at which it was measured is indicated in square brackets.

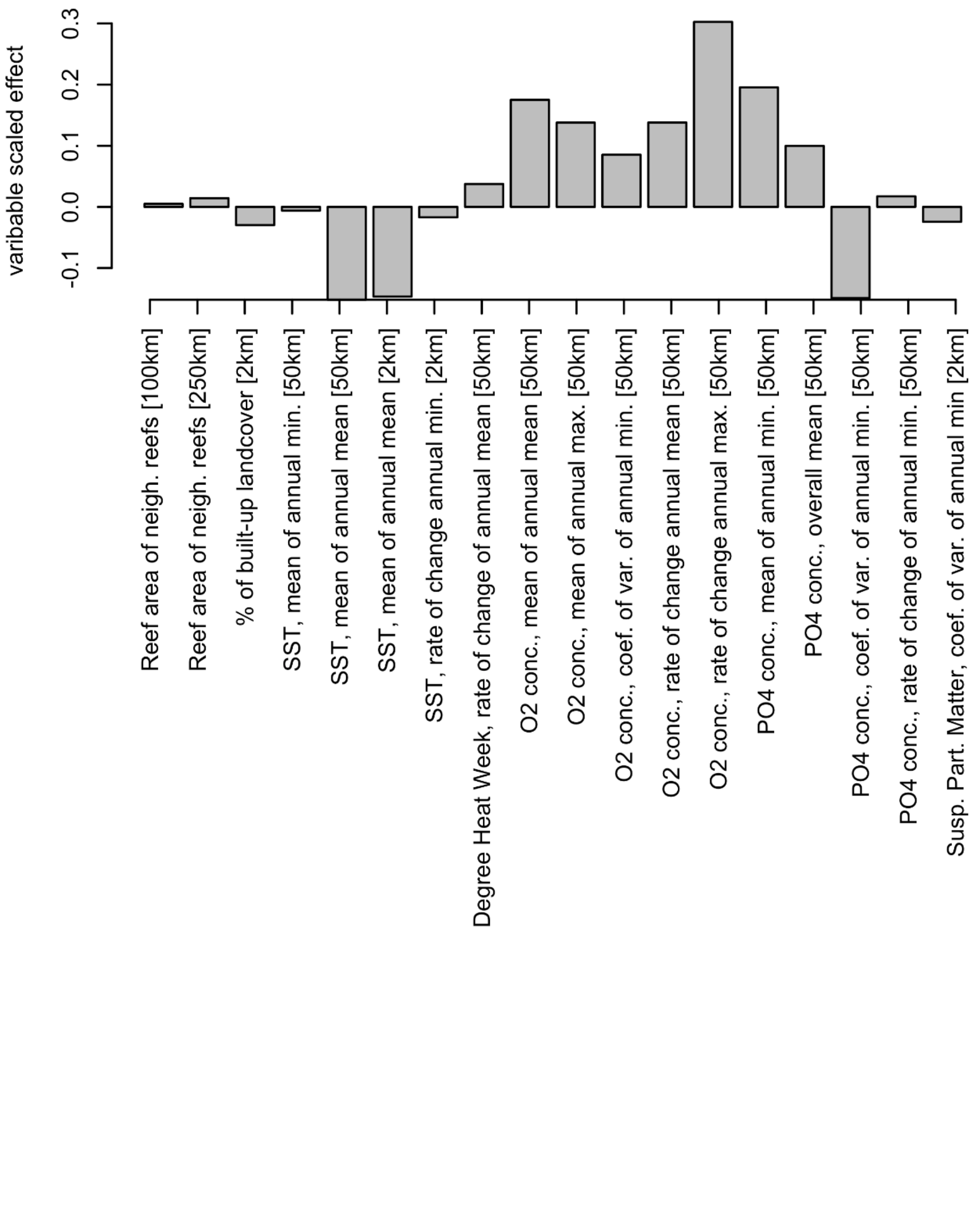

**Supplementary Figure 14. Environmental predictors associated with local effects on genetic distances.**

The panels below show the key environmental predictors retained by the elastic net regularization and stepwise model selection as best explaining local effects on Bray–Curtis genetic distances. For each predictor, the left panel displays its spatial distribution across the world's coral reefs, and the right panel shows its association with the predicted probability of observing a positive local effect on genetic distance, P(ΔBCD > 0). Values close to 1 indicate reefs with predicted positive effects, values near 0 indicate predicted negative effects.

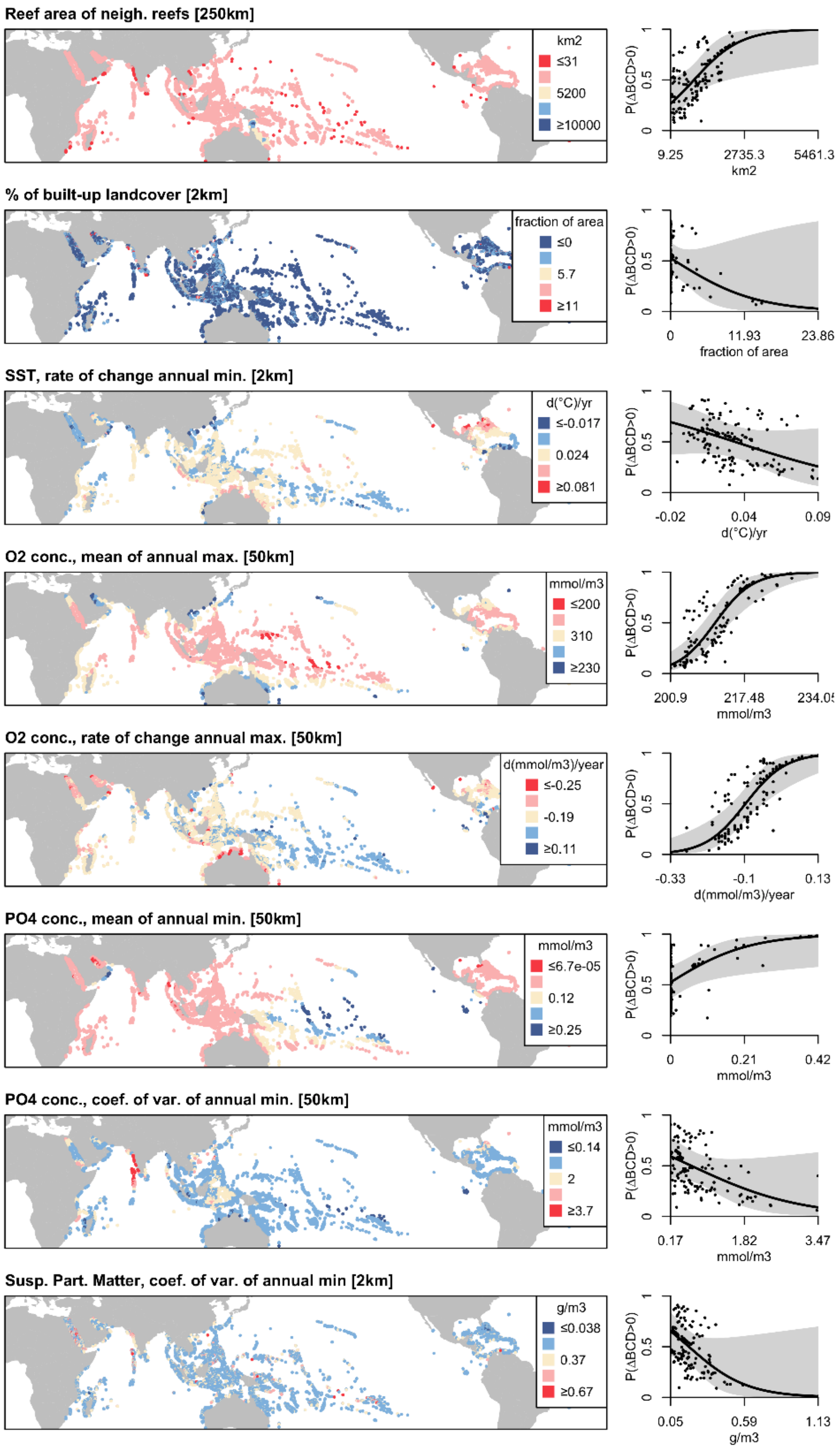
